# Molecular and phenotypic blueprint of the hematopoietic compartment reveals proliferation stress as a driver of age-associated human stem cell dysfunctions

**DOI:** 10.1101/2023.09.15.557553

**Authors:** Emanuele Lettera, Serena Scala, Luca Basso-Ricci, Teresa Tavella, Lucrezia della Volpe, Elena Lo Furno, Kerstin B. Kaufmann, Laura Garcia-Prat, Pamela Quaranta, Raisa Jofra Hernandez, Alex Murison, Kety Giannetti, Alicia G. Aguilar-Navarro, Stefano Beretta, Anastasia Conti, Giacomo Farina, Eugenia Flores-Figueroa, Pietro Conte, Marco Ometti, Ivan Merelli, Stephanie Z. Xie, Alessandro Aiuti, Raffaella Di Micco

## Abstract

Hematopoietic stem/progenitor cell (HSPC) aging studies have been associated with myeloid skewing, reduced clonal output, and impaired regenerative capacity, but quantitative immunophenotypic and functional analysis across human aging is lacking. Here, we provide a comprehensive phenotypic, transcriptional, and functional dissection of human hematopoiesis across the lifespan. Although primitive HSPC numbers were stable during aging, overall cellularity was reduced, especially for erythroid and lymphoid lineages. Notably, HSPC from aged individuals had superior repopulating frequency than younger counterparts in xenografts; yet aged HSPC displayed epigenetic dysregulation of cell cycle, inflammatory signatures, and a reduced capacity to counteract activation-induced proliferative stress with concomitant accumulation of DNA damage and senescence-like features upon xenotransplantation. This phenotype was recapitulated by enforcing proliferative stress *in vivo* on cord blood (CB) HSPC. Overall, our work sheds light on dysregulated responses to activation-induced proliferation underlying HSPC aging and establishes CB xenotransplantation-based models as suitable for studying age-associated hematopoietic defects.

## Introduction

Age-associated changes in the hematopoietic stem and progenitor cell (HSPC) compartment have been extensively unveiled in mice, where a marked myeloid skewing, impaired lymphoid and erythroid potential, and a decreased clonal output of hematopoietic stem cells (HSC) have been reported upon aging^1–3^. Mechanistically, changes in metabolic pathways, compromised maintenance of genome integrity, and epigenetic dysregulation have been linked to rapid exhaustion of the aging stem cell pool and impaired regeneration properties^4–10^. It has also been proposed that the most primitive HSC are predominantly in a quiescent state and may divide only a few times in the lifetime of a mouse^11,12^ to minimize DNA replication-associated mutations, and ultimately stem cell exhaustion. In agreement with this, the proliferative history of HSC has emerged as an important contributor to the dysfunctional output of aged HSC^13–16^. Along the same lines, upon exposure to mitogenic stimuli, murine aged HSC display elevated levels of DNA replication stress associated with cell cycle defects and chromosome gaps or breaks^16,17^. Moreover, as transplantation itself imposes a strong proliferative stress on repopulating HSC, it was reported that transplanted old murine HSC display a significant reduction in long-term repopulating potential compared to young HSC as well as hampered homing and engraftment capacity^7,18^. To date, it is still unclear to what degree the phenotypic observations made in terms of stem cell pool and lineage outputs in mouse settings as well as the underlying molecular principles are conserved during human hematopoietic aging. For instance, the functional decline of the lymphoid compartment, responsible for a defective adaptive immunity in old age, has been observed also in human-aged HSPC leading to the common interpretation that, likewise in mice, human HSPC aging is associated with a prominent myeloid skewing^19–21^. Additionally, the increased incidence of myeloproliferative disorders and hematopoietic myeloid malignancies in aged individuals^22–26^ further supports the above-mentioned shift in myeloid output during aging. Moreover, there is evidence in humans that aging is associated with anemia^27^ and clonal hematopoiesis^23^ which may act as additional cofactors in the development of myelodysplasia and hematopoietic malignancies^28,29^. A growing body of literature has uncovered the functional and molecular heterogeneity of human hematopoietic stem cells throughout life to maintain a constant blood supply under homeostatic steady-state conditions^30–36^. Significant efforts led to the identification of surface markers and gene signatures to unambiguously define the subset of purified HSC responsible for long-term outcomes upon transplantation^32,35,37–40^ and subsequent molecular studies performed in the context of aging had a clear focus on this primitive and rare cell subset. For instance, analysis of proliferation rate in highly purified HSC has revealed altered responses of aged cells to mitogenic stimuli potentially contributing to hematopoietic dysfunctions upon transplant^41,42^. However, it is worth noting that in several therapeutic applications for non-malignant and malignant hematological diseases, a sizable fraction of a heterogenous population of progenitors and long-term HSC needs to be harvested, cultured *ex-vivo* with early-acting cytokines and reinfused in patients to maximize engraftment in the early-phases post-transplant. Furthermore, in the transplantation setting, increasing donor age correlates with higher transplant-related mortality and reduced immune reconstitution^43–45^ while younger donor transplant results in a better long-term outcome with reduced graft-versus-host-disease and increased disease-free survival^46,47^. Thus, characterizing the cellular and molecular landscape of the mature and HSPC compartment at steady-state and understanding the functional impact of aging on HSPC function during *ex-vivo* activation and upon transplantation has the potential to contribute to improving the application of HSPC-based therapies to the elderly.

Here, we report the results of a comprehensive quantitative characterization of blood cell subpopulations in bone marrow and peripheral blood from a large healthy donor cohort across the human lifetime, providing the first comparative analysis of immunophenotypic proportion *vs.* cellular numbers of BM lineages. We combined advanced genomics and *in vivo* transplantation assays to interrogate the functional contribution of proliferative stress to age-related changes in hematopoietic output and stem cell fitness. Overall, our work provides a blueprint of the HSPC compartment upon aging and establishes a relevant transplantation-based model for future studies to further untangle the complex molecular programs underlying human hematopoietic aging and aging-associated diseases.

## Results

### Human aged bone marrow is characterized by a global reduction in hematopoietic output

To dissect the changes in hematopoietic cell composition during aging we first analyzed 45 bone marrow (BM) samples from pediatric (0 to 18 years old, n=8), young adults (18 to 30 years old, n= 10), middle-aged (40 to 60 years old, n= 8), and old (>65 years old, n=19) healthy individuals. As an additional and complementary dataset, we collected 56 peripheral blood (PB) samples from 12 pediatric, 17 young, 8 middle-aged, and 19 old subjects (Supplemental Data Table 1). All the samples were analyzed through our validated multi-parametric flow cytometry protocol^48^ which allows us to unambiguously identify and quantify 25 different hematopoietic subsets, including 10 distinct HSPC subpopulations (See Supplementary Figure 1 for gating strategy). By means of this analysis, we found a significant decrease in bone marrow cellularity during aging in both mature cells and the HSPC compartment (Figure 1A, black outer ticks). Advanced age negatively correlated with monocytes, immature granulocytes (iPMN), dendritic cells (DC), and myeloblasts cell count with the only exception of mature granulocytes (PMN), whose number remained stable over time (Figure 1B-D and Supplementary Figure 2A-B). We also found a gradual decrease of lymphocyte progenitor (Pro-lympho) cell count during aging (Supplementary Figure 2C), while a dramatic drop of B cell production was observed as early as 30 years of age, accompanied by the sharp reduction of B cell Precursors (PreB) and B cell Progenitors (ProB) (Figure 1E-G). BM-resident NK cells showed a stable content over aging, while NKt lymphocytes slightly increased during aged hematopoiesis (Supplementary Figure 2D-E). Similar changes in mature hematopoietic cell composition were also detected in the PB of the analyzed subjects, with marked reduction mainly in the absolute count of B, NK, and T lymphocytes (Supplementary Figure 2H-N). Our analyses also revealed a significant decline in the number of erythroblasts and pro-erythroblasts with age (Figure 1H and Supplementary Figure 2F), in line with the severe anemia reported in elderly subjects^27^.

**Figure 1.**
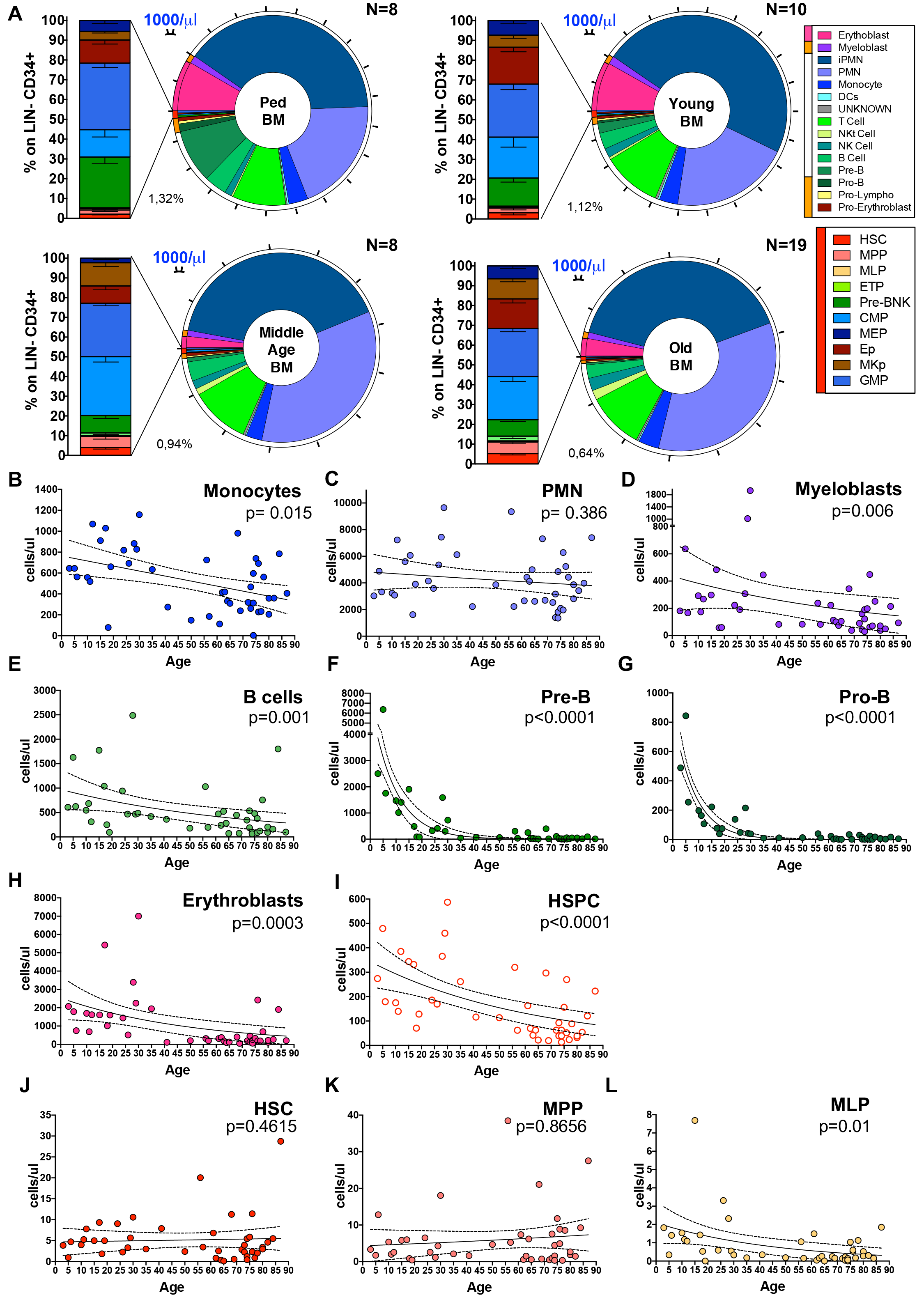
Bone marrow composition of healthy subjects over aging. **A.** (from top left to bottom right) Hematopoietic composition of BM samples from pediatric (< 18 years old, n=8), young adult (< 35 years old; n=10), middle age (<65 years old; n=8) and old (>65 years old; n=19) healthy subjects. Pie charts show the distribution of 25 hematopoietic subsets within the CD45+ gate. Black outer ticks indicate absolute count (1000 cells/µl). Percentages indicate the frequency of the HSPC population (CD34+ LIN-) on total CD45+ cells. The stacked bar graphs indicate the frequency of HSPC subpopulations on LIN-CD34+ cells. **B-L.** Correlation analysis of the absolute counts of each cell subtype (expressed as cells/µL) in the BM with the age of 45 subjects analyzed. HSC, hematopoietic stem cells; MPP, multi-potent progenitors; MLP, multi-lymphoid progenitors; PMN, polymorphonucleated cells.

We next evaluated the changes occurring within the most undifferentiated hematopoietic compartment and found reduced cell count of both CD34+ cells as well as of CD34+LIN-cells (defining the HSPC population) (Figure 1I and Supplementary Figure 2G). Of note, the number of primitive HSC and MPP subsets was stable during aging (Figure 1J-K), while a prominent decline was observed in lymphoid-(MLP and PreB/NK), myeloid-(CMP, GMP) and erythroid (EP and MEP) committed progenitors, except for early T progenitors (ETP) and megakaryocyte progenitors (MKp) which remained stable over aging (Figure 1L and Supplementary Figure 3A-G).

To exclude any potential geographic or collection bias that might account for the observed changes among donors of different age groups, we validated our findings in a North American cohort of 13 donors (age range 21-83y). By comparing HSPC subpopulation frequencies, we found consistent changes in HSPC subsets between the two cohorts of healthy subjects (Supplementary Figure 3H-O). Of note, we observed an increased frequency of primitive subpopulations within the HSPC compartment during aging (Supplementary Figure 3H-I), in line with previous reports^49^, and this increase was also reflected in the higher frequency of phenotypically defined CD49f+ primitive HSC in aged individuals (defined as CD34+CD19-CD38-CD45RA-CD90+CD49f+ cells) (Supplementary Figure 3P).

Altogether, our data indicate that, despite the stable HSC and MPP cell counts, the overall hematopoietic production is severely compromised in aged individuals and that the myeloid skewing reported in previous studies is the result of a more marked reduction of lymphoid cell production (especially in the B cell compartment) with respect to a modest decrease in the myeloid output from the HSPC compartment.

### Human HSPC repopulating frequency is enhanced with age in xenotransplantation assays

The study of aging in human HSC using *in vivo* functional assays is limited, but the evidence suggests that the functional decline of human primitive HSC upon aging is more complex than reported in animal studies^41,42,49–52^. Although we cannot assay the functional activity of human HSC *in vivo* in the bone marrow during steady-state hematopoiesis, transplantation into immunodeficient mice allows us to assess hematopoietic reconstitution potential across aging under the same genetic microenvironment, thus reducing the variability related to extrinsic factors. We undertook a comprehensive limiting dilution xenotransplantation study to quantitatively measure HSPC engraftment of young (21-32y, n=5), middle-aged (52-57y, n=3), and old (78-83y, n=5) BM CD34-enriched cells relative to the neonatal CB-derived CD34+ cells using 314 immunodeficient non-obese diabetic (NOD)-severe combined immunodeficiency (SCID)-IL2Rg-/- (NSG) mice (Figure 2A). Three cell doses (60,000 cells, 10,000 cells, and 1,000 cells) were selected based on a prior study to measure CB functional HSC frequency^40^. Human CD45 engraftment levels were measured at 4-, 12-, and 20-weeks post-transplantation and used to calculate the 1/CD34+ repopulating cell frequency in all four sources of human HSPC (Figure 2B-E, Supplemental Data Table S2, Supplementary Figure 4A). As expected, CB repopulating cell frequency was superior to BM sources, although this was not statistically significant for middle-aged and old BM HSPC at 12 weeks^53^ (Figure 2B, Supplementary Figure 4A). CD34+ repopulating cell frequency kinetics was found to undergo dynamic changes across time with BM sources peaking at 12 weeks and dropping to the lowest frequency at 20 weeks. By contrast, the stem cell frequency for CB was highest at 20 weeks^40^ (1/504 CD34+ cells, Figure 2B, Supplementary Figure 4A). Remarkably, steady-state middle-age and old BM HSPC had a significantly higher CD34+ repopulating cell frequency at 12 weeks and showed superior engraftment relative to young BM HSPC at the 60,000 cell dose for all timepoints (Figure 2C-E). Additionally, at 4 weeks human cell grafts contained lower B cells but a higher fraction of myeloid cells in all the mice transplanted with BM-CD34+ cells with respect to CB, independently from the age of the BM donors likely reflecting a distinct proportion of repopulating progenitors in the two cell sources (Figure 2F,G).

**Figure 2.**
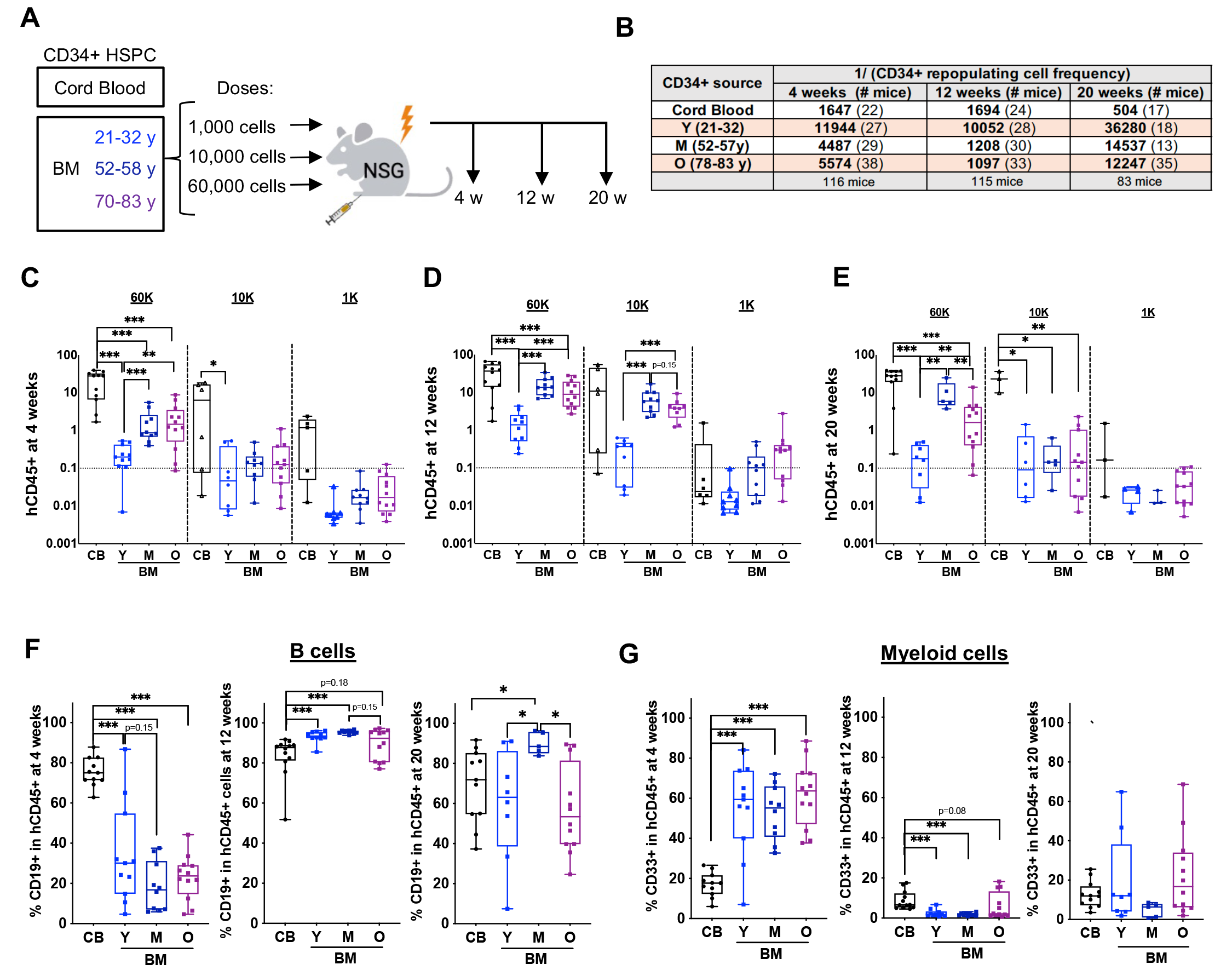
Limiting dilution xenotransplantation assays of uncultured HSPC show a higher frequency of repopulating cells in aged HSPC compared to young HSPC. **A.** Experimental scheme using 3 cell doses in a xenotransplantation time course experiment where human CD45^+^ engraftment and lineage in the bone marrow of NSG mice were assessed at 4, 12, and 20 weeks for the indicated human HSPC cell sources. **B.** The CD34+ repopulating cell frequency for 4, 12 and 20 weeks for CB and BM samples from indicated age groups. **C-E.** Box plots showing human CD45+ cell frequencies in the murine BM in mice transplanted with 60K, 10K and 1K HSPC for CB and BM samples at 4 **(C)**, 12 **(D)**, and 20 **(E)** weeks. **F.** Percentage of CD19+ B cells in the human grafts for mice transplanted with the 60K cell dose at different time points. **G**. Percentage of CD33+ myeloid cells in the human grafts for mice transplanted with the 60K cell dose at different time points. Statistical analyses: Mann-Whitney test.

To understand the molecular underpinnings for the enhanced engraftment of aged human HSPC relative to young, we interrogated the chromatin accessibility landscape of HSPC subpopulations isolated from young and old subjects by ATAC-seq. We found 6 signatures by non-negative matrix factorization (NMF) within 27 HSPC samples, belonging to the distinct HSPC subpopulations (young: 3 donors, 21-28y, n=16; old: 4 donors, 52-79y, n=15; Supplemental Data Table S3, Supplementary Figure 4B). When we assessed the strength of all 6 signatures in young and old CD49f + primitive HSC samples, signature 1 was significantly enriched in old HSC, whilst signature 2 was enriched in young HSC (Supplementary Figure 4C). Transcription factor motif analysis within these two signatures yielded three clusters: shared motifs, motifs enriched in young CD49+ primitive-HSC, and motifs enriched in old CD49+ primitive-HSC (Supplementary Figure 4D and Supplemental Data Table S4 with enrichment data for all motifs identified). Among the shared motifs are well-known HSC regulators including ERG, FLI1, and ETV2^54–56^. Importantly, motifs that are accessible in young CD49+ primitive-HSC are associated with cell cycle regulation (E2F)^57^ and hematopoietic lineage decisions (BCL11a and PU.1)^58^ (Supplementary Figure 4E). In contrast, inflammation response motifs (NFKB and AP1) were enriched in old CD49+ primitive-HSC (Supplementary Figure 4F). Additionally, we found that accessibility of HOXB4 motifs, a factor shown to potently expand HSC^59,60^ was enriched in old HSC (Supplementary Figure 4F).

Together, our data suggest that the enhanced engraftment of aged BM HSPC relative to younger counterparts could be explained by epigenetic alterations in cell cycle, inflammation, and self-renewal factors and prompted us to investigate the biological and functional response of aged HSPC to proliferative stimuli.

### Human aged HSPC show altered proliferation kinetics upon *ex-vivo* activation

We next interrogated the responses of young and old HSPC to a cocktail of human recombinant cytokines and growth factors, including Fms-related tyrosine kinase 3 (Flt3), Stem cell factor (SCF), thrombopoietin (TPO) and IL-3, previously reported to promote proliferation of HSPC without inducing major differentiation^61–63^ and currently representing the gold-standard for HSPC-based gene therapy and transplantation-based clinical applications^64,65^. Specifically, we stimulated BM CD34^+^ cells isolated from young (<35 years old, n=6) and old (>60 years old, n=6) subjects and analyzed their transcriptional profiles, proliferation rates, and clonogenic potential (Figure 3A). Global gene expression analyses identified 1269 statistically significant differentially expressed genes (DEGs) (FDR < 0.05) in CD34+ cells from old subjects compared to their younger counterparts after an overnight activation, of which 529 were upregulated and 740 were downregulated, (Supplementary Figure 5A, Supplemental Data Table S5). Gene set enrichment analysis (GSEA) against hallmark gene sets from Reactome Database revealed upregulation of genes associated to Cell cycle (*RAD17, CDK7, BUB3, CENPN*), Inflammation (*IFIT1, IFIT3, IFIT5, STAT2, OAS2, ISG15, ISG20, SMAD3, BIRC3, TRIM22, CDC23*), Chromatin conformation and Gene expression regulation (*H3-3B, TAF9, HAT1, ELP5, KAT7, SLU7, H2AC6, JUN*), and Protein Translation and localization (*RARS2, EIF4B, WARS1, PEX19, SARS1, TIMM10B, MRSP16*) categories in activated old CD34+ cells with concomitant downregulation of genes belonging to Hematopoiesis (R-HSA-418555; *AVP, PTGIR, OR2V2, ADCY9, GNAZ, GNB2, GRK2, HRH2, GPR176, ASH2L*) and Cellular organization and Transport categories (R-HSA-1474244; *SERPINA1, COPB2, ARCN1, SEC23IP, SCFD1, COPB1, CAPZA1*) (Figure 3B and Supplementary Figure 5B).

**Figure 3.**
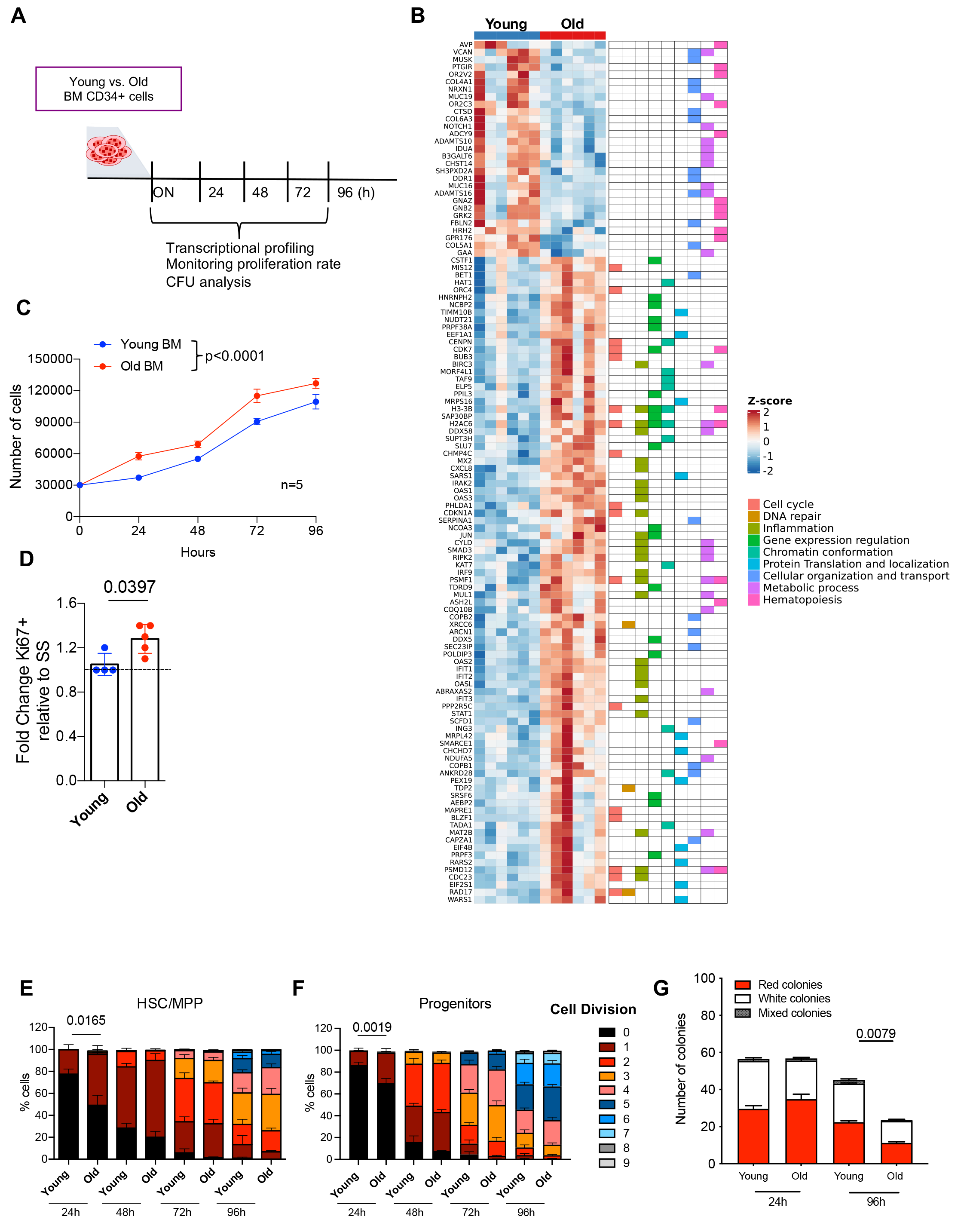
*In vitro* functional characterization of human aged HSPC. **A.** Experimental scheme for *in vitro* proliferation assay. 3×10^5 BM-derived CD34+ cells from young and old individuals were cultured in the presence of the human recombinant cytokines and analyzed at the indicated time points for transcriptome, proliferation rate and clonogenic potential. **B.** Graph showing, on the left, a heatmap with DEGs between young (n=6) and old (n=6) CD34+ cells upon overnight activation in culture and, on the right, the Reactome categories associated with the distinct genes. **C.** Growth curve showing the proliferation of both BM-derived young (n=5) and old (n=5) CD34+ cells. Data are shown as mean± SD. **D.** Fold change of Ki67+ cells relative to overnight activated condition of young (n=4) and aged (n=5) BM-derived CD34+ cells at 24h after culture. Data are shown as mean ± SD. **E-F.** Stacked bar graphs showing the frequencies of cells that completed one or more cell division over time in culture in young and old HSC/MPP (**E**) and Progenitor (**F**) subsets. **G.** Number of erythroid, myeloid and mixed colonies generated starting from 800 BM-derived CD34+ cells of young (n=5) and old(n=5) donors upon activation in culture. Colonies were counted 14 days after plating in CFU-C assay. Data are shown as mean ± SEM. Statistical test for growth curves: Linear mixed-effects model. Statistical test for group comparison: Mann-Whitney test.

Growth curve analysis showed that aged HSPC displayed a higher proliferation rate, mainly early points post-activation, further supported by the higher percentage of Ki67+ proliferating cells as compared to younger counterparts (Figure 3C-D), suggesting a faster exit from quiescence. To better elucidate whether the earlier activation of old compared to young HSPC can be captured in the subset enriched for the most primitive HSPC, we analyzed the kinetics of cell divisions by Cell trace (CT), a fluorescent cytoplasm-binding molecule that allows tracing multiple generations on the basis of dye dilution, on sorted HSC/MPP (defined as LIN-CD34+CD38-CD45RA-cells), and more committed subsets including myeloid, erythroid and megakaryocytes progenitors (defined as LIN+CD34+CD38+CD10-CD7-cells) (Figure 3E-F). Total CD34+ cells and sorted HSPC subsets from both young and old subjects were labelled with CT dye and cultured for 96h to monitor their proliferation rate over time (Figure 3E-F and Supplementary Figure 5C-D). Our assay revealed that a higher percentage of total CD34+ cells as well as of HSC/MPP and committed subsets isolated from old donors completed one or more cell divisions at early time points in comparison to younger counterparts. These early differences in proliferation rate diminished over time with a similar profile of cell divisions observed at later time-points between young and aged HSPC subsets (Figure 3E-F). We next evaluated the clonogenic potential of young and aged CD34+ cells and found that while early upon activation there was a comparable number of colonies, a significant reduction in aged-HSPC derived colonies was observed at later time points (Figure 3G).

Taken together, these data suggest that, after *ex-vivo* activation, old HSPC showed a faster exit from quiescence resulting in altered proliferation rates, characterized by transcriptional changes in cell cycle-related and inflammatory gene categories that may ultimately contribute to reduced clonogenicity upon extended time in culture.

### Activated aged HSPC retain differentiation potential but display impaired long-term fitness after *in vivo* transplantation

To assess the hematopoietic output of activated aged HSPC *in vivo*, we next transplanted at high doses overnight cultured CD34+ cells derived from young (n=6) and old (n=6) donors into immunodeficient NSG mice and compared their human hematopoietic engraftment and blood lineages reconstitution over time (Figure 4A). Upon transplantation, we found no significant differences in the kinetics and the frequency of human CD45+ hematopoietic cells in peripheral blood (Figure 4B), while we observed a reduced human graft in the BM of mice transplanted with CD34+ cells isolated from aged donors at 20 weeks post-transplant (Figure 4B, Supplementary Figure 6A). Of note, we found that mice transplanted with old CD34+ cells showed a statistically significant reduction in both myeloid and B cell output accompanied by a relative increase of T cell content (CD3+) in the peripheral blood at the time of euthanasia (Figure 4C-E). These changes in the lymphoid compartment were also observed in the spleen of the mice transplanted with old CD34+ cells (Figure 4D-E). We also confirmed by quantitative analyses a reduced B cell output both in PB and in BM from mice belonging to the old group accompanied by a reduction of the overall HSPC content at 20 weeks (Figure 4F-G and Supplementary Figure 6B). In line with the differences in HSPC composition observed in old compared to young subjects at steady state (Supplementary Figure 3H-O), we found an increased frequency of LIN-CD34+CD38-cells and a reduced lymphoid progenitor contribution in the human HSPC engrafted in the BM of the mice transplanted with old CD34+ cells (Supplementary Figure 6C-D). In contrast, absolute quantification of distinct HSPC subsets in the BM of the two groups of transplanted mice unveiled a reduced amount of primitive but not more committed HSPC progenitors in the old group (Supplementary Figure 6E). Moreover, CD34+ cells retrieved at the endpoint from mice transplanted with old CD34+ cells showed reduced clonogenic potential with respect to their younger counterparts when rechallenged in colony assays (Figure 4H). Interestingly, gene expression analyses on CD34+ cells isolated at sacrifice from transplanted mice belonging to the old group showed increased levels of cell cycle inhibitor *p21* and several pro-inflammatory cytokines (including *IL6*, *IL8*, *IL1β, MCP1*, and *TNFα*) compared to younger counterparts (Figure 4I-N), together with the accumulation of DNA double-strand breaks (DSB), detected through the phosphorylation of histone H2A.X (γH2AX) (Figure 4O).

**Figure 4.**
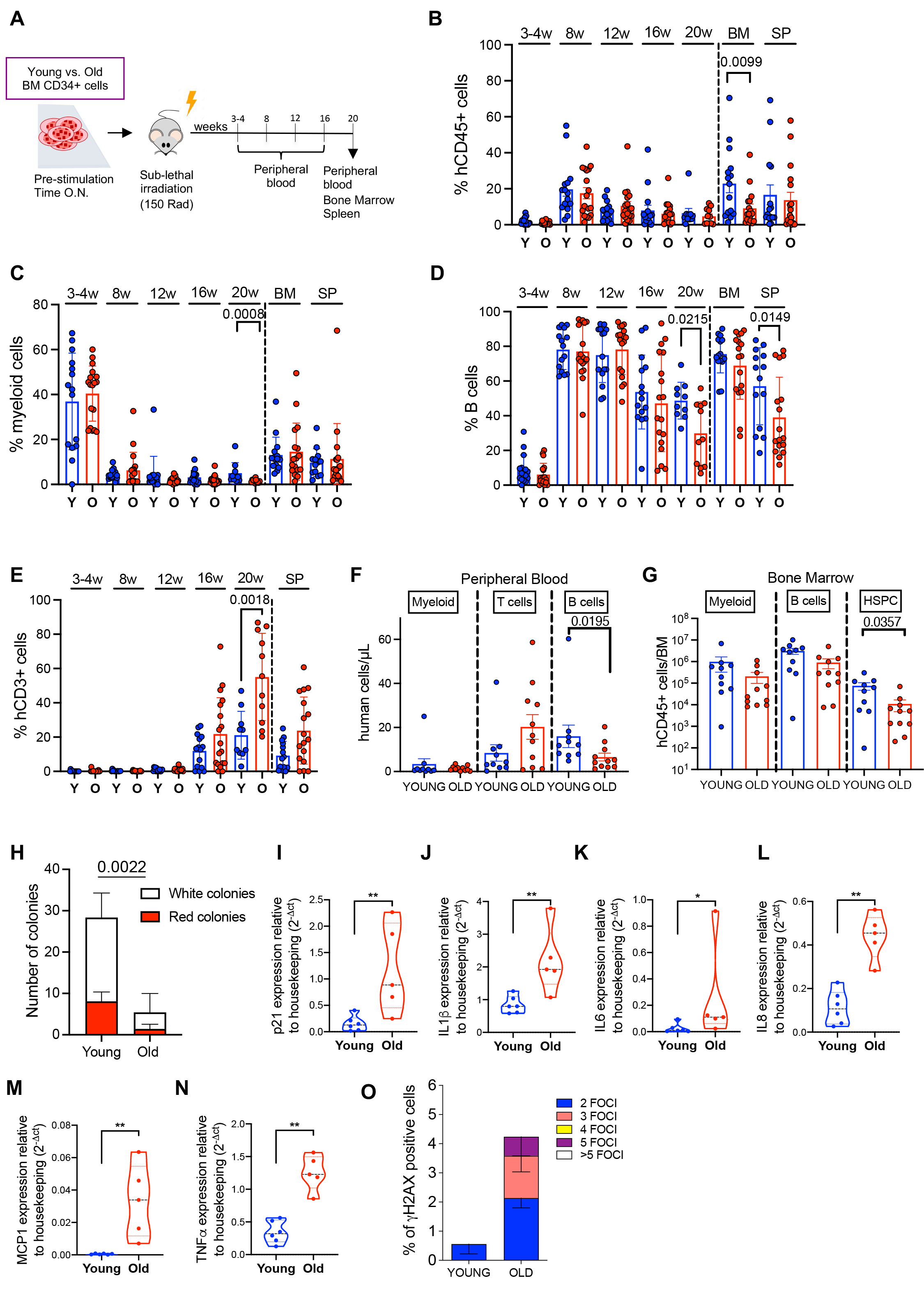
*In vivo* functional characterization of pre-activated human aged HSPC. **A.** Percentage of human engrafted cells (CD45+ cells) in PB overtime as well as in BM and Spleen (SP) at euthanasia in mice transplanted with young or aged HSPCs. Young donors n=4, Old donors n=4, (n=3 for each donor)**. B-D.** Percentage of human myeloid cells (CD13+ cells), B cells (CD19+ cells) and T cells (CD3+ cells) in PB overtime as well as BM and SP at 20 weeks in mice transplanted with young or old HSPCs. **E-G.** Absolute quantification of hematopoietic populations in PB and BM at 20 weeks after transplantation in mice transplanted with young and old HSPCs. **H-N.** Relative mRNA expression of cell cycle inhibitor *p21* and inflammatory genes *IL1β, IL6, IL8, MCP1* and *TNFα* in human CD34+ cells isolated from BM of mice transplanted with young and aged HSPCs (20 weeks). Gene expression was measured by quantitative Real-Time PCR and represented as 2229 cl:1029-ΔCt relative to housekeeping Statistical test: Mann-Whitney test. **O.** Frequencies of cells showing nuclear ψH2AX foci in human CD34+ cells isolated from BM of mice transplanted with young and old HSPCs at 20 weeks.

These data suggest that, although activated aged HSPC reconstitute all main blood cell lineages (myeloid, T, and B cells) shortly upon transplantation, they are likely unable to counteract transplant-related proliferation stress resulting in detrimental cellular responses that impair their functionality in the long-term.

### Establishing an *in vivo* model of proliferation stress by low cell dose transplantation

To study the impact of proliferative stress on hematopoietic reconstitution *in vivo,* we hypothesized that, when transplanted in limited numbers, HSPC may undergo an increased number of cell divisions to repopulate the bone marrow than when transplanted in higher amounts^40^. We therefore transplanted overnight cultured neonatal CB-derived HSPCs at different cell doses in sub-lethally irradiated immune-compromised mice and tested their repopulating capacity and blood lineage reconstitution at early (6 weeks) and late (20 weeks) phases after transplantation. We designed three experimental groups: a high-dose group (5×10^4^ CD34+ cells), an intermediate-dose group (1×10^4^ CD34+ cells), and a low-dose group (3×10^3^ CD34+ cells) (Figure 5A). Concomitantly, to validate that low engraftment in transplantation experiments associates with a higher proliferation rate of the infused cells, we combined cell number measurements with a previously established short-timescale dual pulse labelling method^66^ that allows us to quantify the number of cells entering S-phase per hour. Specifically, before euthanasia, NSG mice were subjected to an initial intravenous EdU pulse (2 hours) followed by a second intravenous injection of BrdU. By measuring the cells positive for one or both markers we estimated the number of cells in active proliferation at early and late phases after transplantation.

**Figure 5.**
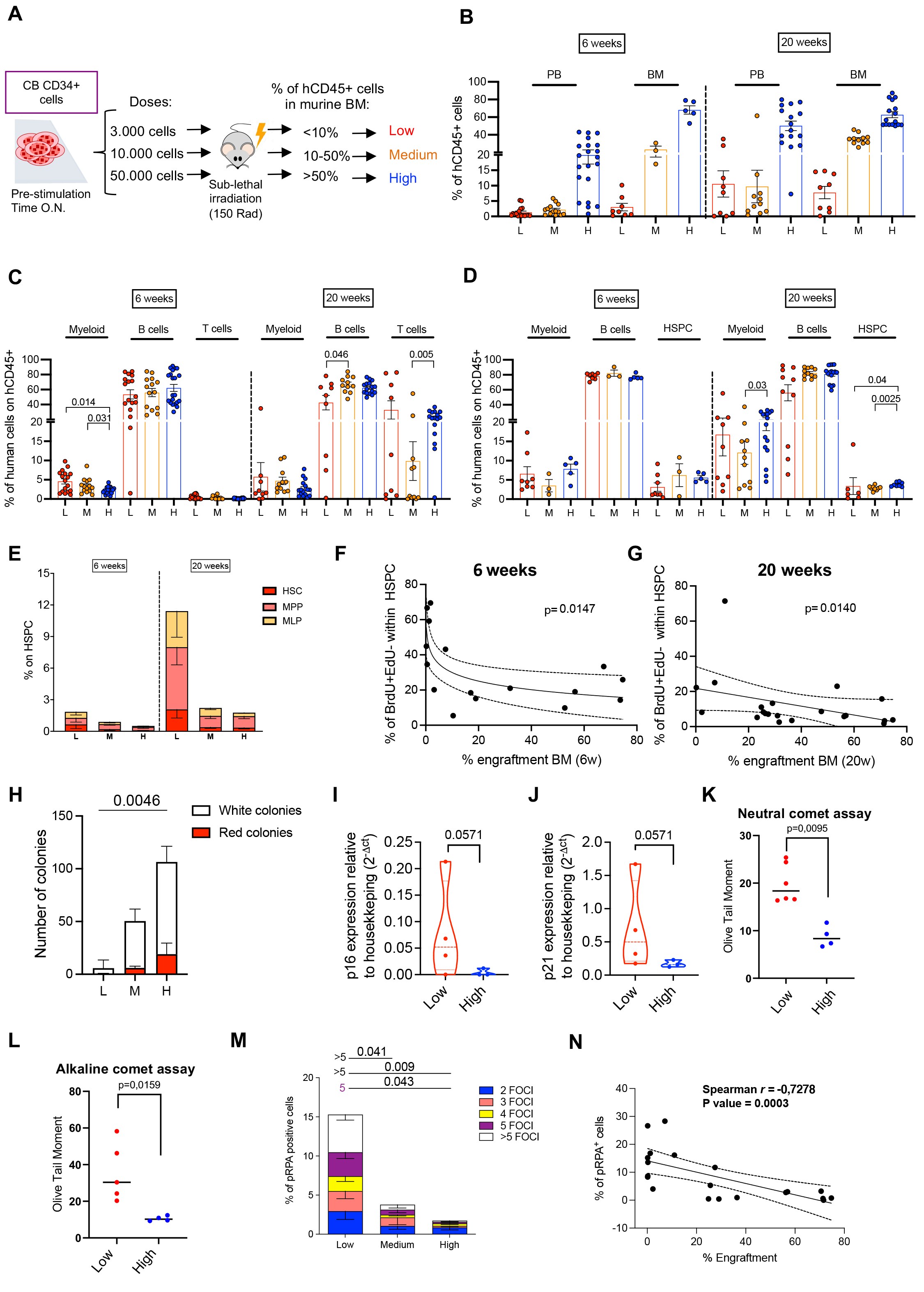
*In vivo* modeling of proliferation stress. **A.** Schematic representation of the *in vivo* transplantation setting. Three groups of mice are transplanted with different doses of CB-derived CD34+ cells. After evaluation of human cell content in the murine BM at euthanasia, mice were divided into 3 distinct categories according to the level of engraftment: Low-engrafted (L, <10% hCD45+ cells), Middle-engrafted (M, 10-50% of hCD45+ cells) and High-engrafted (H, >50% hCD45+ cells) **B.** Percentage of human engrafted cells (CD45+ cells) in PB and BM of transplanted mice at both 6 and 20 weeks after transplantation**. C.** Percentage of human myeloid cells (CD33+CD66b+ cells), B cells (CD19+ cells), and T cells (CD3+ cells) in PB of transplanted mice at both 6 and 20 weeks after transplantation. **D.** Percentage of human myeloid cells (CD33+CD66b+ cells), B cells (CD19+ cells), and HSPC (CD34+ cells) in BM of transplanted mice at both 6 and 20 weeks after transplantation. **E.** Percentage of primitive HSPC subsets within BM HSPC compartment in transplanted mice at both 6 and 20 weeks after transplantation. **F-G.** Correlation between the percentage of HSPCs that entered S phase per hour (BrdU+EdU-) with human engraftment in murine BM at 6 and 20 weeks after transplantation. **H.** Number of colonies generated starting from the same amount of CD34+ cells isolated from transplanted mice at 20 weeks. **I-J.** Relative mRNA expression of cell cycle inhibitors *p16* and *p21* in human CD34+ cells isolated from low or high-engrafted mice (20 weeks). **K-L.** Olive Tail Moment average values of neutral (**K**) or alkaline (**L**) comet experiments conducted in CD34+ isolated at 20 weeks after transplantation from Low or High engrafted mice. **M.** Frequencies of cells showing nuclear pRPA foci in human CD34+ cells isolated from BM of transplanted mice (20 weeks). Statistical comparison test: Kruskal-Wallis with Dunn’s correction for multiple comparisons. Only statistically significant p-values were reported. **N.** Correlation between the frequency of pRPA+ cells within HSPC with human engraftment in murine BM at 20 weeks after transplantation.

To assess the effect of proliferation stress on the engraftment, we further stratified our transplanted mice according to both the number of cells infused and their engraftment in the bone marrow (Figure 5B; low-engrafted: <10% of hCD45+ cells; middle-engrafted: 10-50% of hCD45+ cells; high-engrafted: >50% of hCD45+ cells). Interestingly, we observed that low-engrafted mice displayed a phenotype of hematopoietic reconstitution like that observed after transplantation of old CD34+ cells in NSG mice. Indeed, we found slightly increased myeloid output at 6 weeks in the PB of low-engrafted mice and reduced B cell production with a tendency of increased T cell output (4 out of 9 mice) in the PB at 20 weeks (Figure 5C). At the same time point, low-engrafted mice showed a significant decrease in the percentage of human HSPC in the BM, with an expansion of more primitive cells (defined as LIN-CD34+CD38-) and a concomitant decrease in the percentage of B lymphoid progenitors (PreB/NK), a phenotype also observed in mice transplanted with old CD34+ cells (Figure 5D-E and Supplementary Figure 7A). Moreover, the evaluation of EdU and BrdU uptake revealed a statistically significant negative correlation between the percentage of HSPC entering in S phase per hour (defined as EdU-BrdU+ cells)^66^ or total proliferating HSPC (EdU-BrdU+, EdU+BrdU+, and EdU+BrdU-cells) and the human graft in the BM at 6 weeks and 20 weeks post-transplantation (Figure 5F-G and Supplementary Figure 7B-C). In addition, we also observed that low-cell engraftment was associated with higher T cell proliferation, suggesting that the increased T cell frequencies observed in the low-engrafting mice at 20 weeks was likely due to mature T cell proliferation in the PB rather than *de novo* T cell production (Supplementary Figure 7D).

Finally, we reasoned that the increased proliferation rate imposed by low-cell transplantation could lead to premature exhaustion of HSPC, thus recapitulating the defects observed upon activation in transplanted aged HSPCs. Strikingly, when plated at the same number, we found a reduced clonogenic capacity of CD34+ cells retrieved at the time of euthanasia (20 weeks post-transplantation) from the BM of low-engrafted mice with respect to middle/high-engrafted mice (Figure 5H). Moreover, CD34+ cells isolated from mice transplanted with low cell doses displayed a significant increase in expression levels of senescence-associated cell cycle inhibitors *p16* and *p21* (Figure5I-J) as well as in inflammatory genes *IL1-α*, *MCP-1*, and *TNF-α* (Supplementary Figure 7E-G). Moreover, the proliferative stress imposed by the low-cell dose was associated also with the higher presence of both single- and double-stranded DNA breaks, measured through comet assay^67^ (Figure 5K-L and Supplementary Figure 7H-I). Finally, CD34+ cells isolated from low-engrafting mice showed a statistically significant increased frequency of pRPA-positive cells (Figure 5M), a well-established marker of DNA replicative stress^68^ which inversely correlated with the engraftment levels in the bone marrow (Figure 5N).

Taken together, these findings indicate that the proliferative burst imposed by low cell transplantation in overnight cultured HSPC recapitulates age-associated changes of human hematopoiesis and leads to the activation of detrimental cellular responses characterized by DNA damage accumulation and inflammatory programs, ultimately leading to altered hematopoietic output and premature exhaustion of HSPCs.

## Discussion

Age-associated changes in the hematopoietic compartment include relative expansion of functionally defective HSC, skewed myeloid differentiation at the expense of T/B lymphocytes, and loss of stem cell clonality. However, this knowledge was mainly inferred from murine studies and the translation of these findings to the human context has yet to be defined, given the different life spans of rodents with respect to primates. Here, we provided a comprehensive quantitative characterization of the changes occurring during hematopoietic aging in a cohort of 74 healthy donors, comprising PB and BM collected from pediatric as well as young, middle-aged, and old subjects. By virtue of quantitative measurements of the overall hematopoietic system, we found that human aged hematopoiesis at steady state is characterized by an overall reduction of differentiated cell production, primarily impacting lymphoid and erythroid lineages, accompanied by a more modest decrease of myeloid populations. Thus, we reason that the reported myeloid skewing observed in previous works^49^ might be the result of the distinct decreasing trend in the myeloid *vs*. lymphoid/erythroid compartments. In addition, by comparing the analysis of immunophenotypic cellular numbers *vs.* the proportion of hematopoietic output, we report the previously unappreciated observation that primitive human HSC and MPP populations display constant numbers across lifetime. Therefore, there must be compensatory mechanisms acting at the HSC level to counteract their reduced functionality to support hematopoietic homeostasis. Intriguingly, our *in vivo* xenograft data of aged HSPC showed that human-aged HSPC are not intrinsically impaired in the capacity to differentiate into lymphoid lineages. Indeed, upon xenotransplantation, aged CD34+ cells were able to generate *de novo* B and T cells, suggesting that extrinsic signals in the host microenvironment may drive HSPC differentiation towards specific lineages. In support of this hypothesis, it was described that inflammatory cytokines derived from non-hematopoietic cell types generate an age-associated chronic inflammation, also known as “inflammaging”^69,70^, that might lead to reduced lymphoid production. This “subclinical” inflammatory environment might also affect erythroid differentiation. Indeed, in line with the known age-associated anemia^27,71^ our data showed that the entire erythroid lineage is reduced during aging, with a progressive decrease of both immature and committed populations. Since previous studies suggested a reduced sensitivity to erythropoietin (EPO) of aged HSPC^71^ further investigations will be required to assess intrinsic *vs.* extrinsic contribution to this phenomenon.

Strikingly, our *in vivo* xenograft data of aged HSPC transplanted at steady state showed that human-aged HSPC display a superior repopulating frequency than younger counterparts, in agreement with another study where CD34+CD38-CD90+CD45RA-HSC cells from aged BM donors display higher engraftment compared to young BM donor samples^42^. Using a novel inflammation-recovery xenotransplantation model some of us recently discovered an inflammatory memory HSC (HSC-iM) subset that is equivalent to a subset of HSC found in the bone marrow of old adults^94^. HSC-iM retain transcriptional and epigenetic memory of prior inflammation concordant with signatures associated with T cell memory and inflammatory memory in epithelial stem cells (e.g. AP-1 transcriptional and epigenetic upregulation). Accordingly, many of the top motifs enriched in the chromatin accessibility signature from aged *vs.* young immunophenotypic CD49f+ primitive HSC (AP-1 and NFkB) at steady-state are the same motifs enriched in HSC-iM after recovery from inflammatory challenge^72^. Considering these findings, we hypothesize that repeated inflammatory stress events in the form of acute infections across the human lifespan are an “allostatic load” that impacts HSC function^73^. Indeed, our xenograft data are reminiscent of the engraftment patterns observed in serial transplants after CB HSPC were challenged with lipopolysaccharide (LPS) to mimic bacterial infection, where the repopulating frequency of LPS-treated CB was significantly higher than untreated controls in quantitative secondary transplants^94^.

Moreover, this inflammatory memory of aged HSPC may sensitize them to damage, heightening their response to the proliferative stress induced by *ex-vivo* activation and transplantation, as previously reported for skin epithelial cells and akin to trained immunity in HSC^74–79^. Indeed, when activated, aged HSPC displayed a higher proliferation rate with a more rapid exit from dormancy associated with accumulation of proliferation stress-induced DNA damage, cell cycle inhibitors, inflammatory cytokines, and reduced long-term engraftment. Altogether, these findings clearly suggest that aging alters the transcriptional and epigenetic state of human HSPC resulting in functional defects upon activation. The enhanced ability of aged HSPC, and of sorted HSC/MPP subset, to respond to mitogenic stimuli *ex-vivo* appear in contrast with the recent evidence reporting that individual CD49f^+^ primitive HSC showed age-associated delayed timing of first division and prolonged G1 phase^41^ and likely highlight a differential response to mitogenic stimuli between a pool of progenitors/stem cells compared with highly purified primitive cells. Instead, our findings align with literature reporting increased DNA replication stress in aged murine HSC during the S phase^16^ and with the evidence that proliferation triggered by simulated viral infection leads to single-strand DNA break accumulation in murine HSC^80^. As kinetics of G0/G1 transition are independently regulated in human HSC^40^ and differ across stem cell subsets it is worth speculating that multiple points of cell cycle progression may be affected in aged HSPC. This is concordant with the possibility that alternative states of quiescence in human HSC are disrupted during aging; such that aging has shifted the HSC pool to a pool of short-term repopulating cells that give rise to rapid hematopoietic reconstitution following transplantation by progressively depleting the pool of latent HSC with long-term repopulating potential^36^.

*Ex vivo* activation in response to mitogenic stimuli has been also shown to affect additional HSC quality-control programs including mitochondrial metabolism, protein synthesis, autophagic recycling and UPR stress response early upon exposure to culture systems and prior to cell cycle progression^32,81–84^. As we show that a stimulation time of approximately 16h is sufficient to affect the long-term repopulating potential of aged HSPC in transplantation experiments, one can speculate that the observed loss of aged HSC fitness may result from the combinatorial effects of dysregulated culture adaptation programs and the proliferative stress imposed by the transplant.

Importantly, the altered aged phenotype observed in mice transplanted with aged CD34+ cells was recapitulated in mice transplanted with a low input of pre-activated neonatal CB cells. The higher proliferation rate during the early stage of blood reconstitution observed in mice was associated with the accumulation of DNA damage, the establishment of an inflammatory program, and exhaustion of transplanted HSPC at later stages. From a clinical perspective, our *in vivo* proliferation-stress model can be also used to test innovative strategies to mitigate stress-induced defects of human hematopoiesis. Indeed, apart from the impaired HSPC functionality during physiological aging, it was also reported that the HSCT-induced proliferation of HSPC, due to the need to repopulate the host BM and reconstitute the hematopoietic system, might be responsible for the reduction of HSPC clonality in transplanted patients in the long-term^85^. These effects might also be exacerbated in the case of limited HSPC graft size due to patients’ intensive pre-treatments or other underlying conditions^65^. Finally, even if most autologous gene therapy treatments occur in pediatric patients or young adults^64,65^, applications to cancer and degenerative diseases are predicted to increase in the next decades. In this setting, hematopoietic stress and inflammation could be further amplified upon *ex vivo* culture of autologous stem cells from old subjects leading to reduced long-term functionality and altered hematopoietic reconstitution, possibly affecting both the health span and lifespan of treated patients. Altogether, our data show that human hematopoietic aging is associated with overall hematopoietic reduction with impaired functionality and altered proliferation-stress responses acting at the HSC level. As the above-mentioned xenotransplantation-based models of neonatal HSC greatly mirror the dysfunctions observed in repopulating aged HSPC, they may represent a powerful tool for understanding the mechanisms driving aging of human HSPC with clinically relevant implications.

## Methods

### Human hematopoietic samples in the Italian Cohort

We complied with all the ethical regulations for retrieving biological materials from healthy donors. To characterize the hematopoietic compartment over aging we collected BM samples from 8 pediatric (0 to 18 years old), 10 young adults (18 to 30 years old), 8 middle-aged (40 to 60 years old) and 19 aged (>65 years old) healthy individuals as well as PB samples from 12 pediatric (0 to 18 years old), 17 young adults (18 to 30 years old), 7 middle-aged (40 to 60 years old) and 14 aged (>65 years old) healthy subjects. Pediatric BM samples were collected as residual excess material from subjects undergoing BM harvest as donors for transplantation, after parents’ signing informed consent. Aged BM samples were collected from subjects who underwent hip replacement surgery at San Raffaele Hospital, Milan, after obtaining written informed consent. BM CD34+ cells from young healthy donors for functional assays were collected at San Raffaele Hospital or purchased from Lonza. For the phenotypic analyses the sample size was determined by the number of individuals for whom excess material was available, after signing informed consent for research protocols approved by the San Raffaele Scientific Institute’s Ethics Committee (according to protocol TIGET09). We excluded from our analyses all subjects with compromised immune functions and previous history of cancer or cancer treatments. We also excluded subjects that were positive to HBV, HCV, and HIV infections. We included only subjects that scored as “no frail” upon Edmonton Frailty Scale (EFS)^86^ evaluation.

### Human hematopoietic samples in the North American Cohort

Human bone marrow mononuclear cells (MNC) were either purchased from Lonza, collected from young healthy volunteers from Toronto with informed consent, or collected from patients undergoing hip replacement surgery at the at the Traumatology and Orthopedics Hospital Lomas Verdes (IMSS), Mexico with verbal consent, as determined by the Institutional Ethical Board. The Mexican samples were confirmed to have no dysplasia of any hematopoietic lineages by histological and complete blood count analysis^87^. Ethical approval was obtained from the Institutional Review Board (R-2012-785-092). Samples were viably frozen and stored at -150°C. All Toronto hematopoietic samples were collected with informed consent according to the procedures approved by the University Health Network (UHN) Research Ethics Board (REB 01-0573-C). CB samples were obtained from Trillium and Credit Valley Hospital and William Osler Health Centre, processed as previously described, and stored viably as CD34 enriched HSPC at -150°C^84^. BM samples were thawed and CD34 enrichment from BM samples was performed by positive selection with the CD34 Microbead kit (Miltenyi) with MACS magnet technology (Miltenyi) and stained with the following panel for HSPC immunophenotypic analysis: FITC-anti-CD45RA, PE-anti-CD90, APC-anti-CD10, PE-Cy5-anti-CD49f, BV711-anti-CD19, APC-Cy7-anti-CD34, PE-Cy7-anti-CD38, biotin-anti-CD135, Alexa-Fluor-anti-CD7, V421-anti-CD33, BV650-anti-CD71, streptavidin-Qdot605, and propidium iodide.

### Whole Blood Dissection Staining

PB samples and BM aspirate were stained according to the published Whole Blood Dissection (WBD) protocol^48^. In brief, after red-blood cell lysis, the samples were labeled with the following fluorescent antibodies:

- Mouse anti-human CD3-BV605 (Clone: OKT3; Biolegend, 317322), Verified Reactivity: Human; Application: Flow cytometric analysis of antibody surface-stained cells. Dilution: 1:50
- Mouse anti-human CD56-PC5 (Clone: 5.1H11; Biolegend, 362516), Verified Reactivity: Human; Application: Flow cytometric analysis of antibody surface-stained cells. Dilution: 1:50
- Mouse anti-human CD14-BV510 (Clone: M5E2; Biolegend, 301842), Verified Reactivity: Human, Cynomolgus, Rhesus; Application: Flow cytometric analysis of antibody surface-stained cells. Dilution: 1:50
- Mouse anti-human CD33-BB515 (Clone: WM53; BD Biosciences, 564588), Verified Reactivity: Human (QC Testing); Application: Flow cytometry (Routinely Tested). Dilution: 1:50
- Mouse anti-human CD41/CD61-PC7 (Clone: A2A9/6; Biolegend, 359812), Verified Reactivity: Human; Application: Flow cytometric analysis of antibody surface-stained cells. Dilution: 1:50
- Mouse anti-human CD66b-BB515 (Clone: G10F5; BD Biosciences, 564679), Verified Reactivity: Human (QC Testing); Application: Flow cytometry (Routinely Tested). Dilution: 1:50
- Mouse anti-human CD7-BB700 (Clone: M-T701; BD Biosciences, 566488), Verified Reactivity: Human (QC Testing), Rhesus, Cynomolgus, Baboon (Reported); Application: Flow cytometry (Routinely Tested). Dilution: 1:50
- Mouse anti-human CD45-BUV395 (Clone: HI30; BD Biosciences, 563792), Verified Reactivity: Human (QC Testing); Application: Flow cytometry (Routinely Tested). Dilution: 1:33
- Mouse anti-human CD38-BUV737 (Clone: HB7; BD Biosciences, 612824), Verified Reactivity: Human (QC Testing); Application: Flow cytometry (Routinely Tested). Dilution: 1:33
- Mouse anti-human CD90-APC (Clone: 5E10; BD Biosciences, 559869), Verified Reactivity: Human (QC Testing), Rhesus, Cynomolgus, Baboon, Pig, Dog (Tested in Development); Application: Flow cytometry (Routinely Tested). Dilution: 1:33
- Mouse anti-human CD135-PE (Clone: BV10A4H2; Biolegend, 313306), Verified Reactivity: Human; Application: Flow cytometric analysis of antibody surface-stained cells. Dilution: 1:33
- Mouse anti-human CD11c-BV650 (Clone: B-ly6; BD Biosciences, 563404), Verified Reactivity: Human (QC Testing); Application: Flow cytometry (Routinely Tested). Dilution: 1:20
- Mouse anti-human CD10-BV786 (Clone: HI10a; BD Biosciences, 564960), Verified Reactivity: Human (QC Testing), Rhesus, Cynomolgus, Baboon (Tested in Development); Application: Flow cytometry (Routinely Tested). Dilution: 1:20
- Mouse anti-human CD34-BV421 (Clone: 561; Biolegend, 343610), Verified Reactivity: Human; Application: Flow cytometric analysis of antibody surface-stained cells. Dilution: 1:20
- Mouse anti-human CD45RA-APCH7 (Clone: HI100; Biolegend, 304128), Verified Reactivity: Human; Reported Reactivity: Chimpanzee; Application: Flow cytometric analysis of antibody surface-stained cells. Dilution: 1:20
- Mouse anti-human CD71-BV711 (Clone: M-A712; BD Biosciences, 563767), Verified Reactivity: Human (QC Testing); Application: Flow cytometry (Routinely Tested). Dilution: 1:20
- Mouse anti-human CD19-APCR700 (Clone: SJ25C1; BD Biosciences, 659121), Verified Reactivity: Human; Application: Flow cytometry. Dilution: 1:20

All the antibodies were purchased from Biolegend and BD Biosciences and they are well characterized and validated by providers. Titration assays were performed to assess the best antibody concentration. After surface marking, the cells were incubated with PI (Biolegend) to stain dead cells. Absolute cell quantification was performed by adding Flowcount beads (BD Biosciences) to samples before WBD procedure. All stained samples were acquired through BD LSR-Fortessa (BD Biosciences) cytofluorimeter after Rainbow beads (Spherotech) calibration. Raw data were collected through DIVA software Version 8.0.2 and analyzed with FlowJo software Version 10.5.3(BD Biosciences). Technically validated results were always included in the analyses, and we did not apply any exclusion criteria for outliers.

### Limiting dilution xenotransplantation study of uncultured human BM HSPC across aging

All animal experiments were done in accordance with institutional guidelines approved by the University Health Network Animal care committee. Aged match female NSG mice (NOD.Cg *Prkdc*scid*Il2rg*tm1Wjl /SzJ; Jackson Laboratory) 10-12 weeks of age were sublethally irradiated with 225 rads 1 day before intrafemoral injection. CB CD34-enriched cells and BM samples were thawed, and CD34 enrichment from BM samples was performed by positive selection with the CD34 Microbead kit (Miltenyi) with MACS magnet technology (Miltenyi) the morning of xenotransplantation. A limiting dilution assay (LDA) to quantitatively measure CD34 repopulating frequency across aging (CB, young BM (21-32y, n=5), middle-age BM (52-57y, n=3), and old BM (78-83y, n=5) in 314 NSG mice was designed based on a prior study to measure CB functional HSC frequency in NSG^40^. CD34-enriched samples were injected into the right femur of mice at three cell doses (60,000 cells, 10,000 cells, and 1,000 cells). Flow cytometry analysis was conducted on an aliquot of each sample to quantify the number of CD34+ cells injected using the following panel: PE-anti-CD19, APC-anti-CD34, PECy7-anti-CD38, FITC-anti-CD45RA and propidium iodide. Each sample was distributed across the indicated cell doses for transplant (n=3-11 mice/cell dose) based on available sample material (see Supplemental Data Table 1). Human engraftment in the bones of mice were assessed at 4, 12, and 20 weeks post-transplantation as previously described by flow cytometry analysis on a BD Celesta with the following antibodies: PE-Cy5-anti-CD45, V500-anti-CD45, PE-anti-GlyA, PE-anti-CD19, BV786-anti-CD33, APC-anti-CD34, FITC-anti-CD71, APC-Cy7-anti-CD41, and BV605-anti-CD56 and a viability dye^40,84^ (sytox blue, ThermoFisher). Mice were considered engrafted if CD45+ cells were >0.05%. CD34 repopulating frequency was estimated using ELDA software^88^ (http://bioinf.wehi.edu.au/software/elda/).

### ATAC-Seq library preparation and analysis

Library preparation for ATAC-Seq was performed on 1000-2000 cells with Nextera DNA Sample Preparation kit (Illumina), according to previously reported protocol^89^. 4 ATAC-seq libraries were sequenced per lane in HiSeq 2500 System (Illumina) to generate paired-end 50-bp reads. Raw sequencing reads were aligned to the hg38 genome build using BWA^90^. All duplicate reads, and reads mapped to mitochondria, chrY, an ENCODE blacklisted region or an unspecified contig were removed (ENCODE Project Consortium, 2012). The set of samples (n=31 subfractions, n=3 young donors (21-28y), n=4 mid/old donors (52-79y)) that passed quality control and used for downstream analysis is in Supplemental Table 3. The catalog of all peaks called in any population was produced by merging all called peaks which overlapped by at least one base pair using bedtools, and divided into 500bp windows. The MACS bdgcmp function^91^ was used to compute the fold enrichment over background for all populations, and the bedtools map function was used to identify the maximum fold enrichment observed at each window covering a peak in the catalog in each population. Maximum fold enrichment at each site in the catalog was quantile normalized between samples, we define this as the ATAC-Seq signal for further analyses. The NMF package^92^ was used to perform non-negative matrix factorization on the quantile normalized fold enrichment at each peak in the catalog (using the brunet method with default parameters) for between 3 and 10 signatures. 6 signatures was selected as optimal. or each signature, sites were ranked according to the strength of each signature, and the top 5% of sites identified as being the sites for each signature. Homer^93^ was used to identify uniquely enriched motifs within the top 5% of sites for each signature, using default parameters. Contiguous genomic windows were merged and the catalog of all called peaks was used as a background. Sites enriched in the top 5% of sites of both signatures were removed prior to computing enrichment.

### Total RNA-seq library preparation and analysis

Whole transcriptomic analysis was performed on a pool of BM CD34+ cells derived from three young and old donors. All conditions were performed in triplicate. Total RNA was isolated after an overnight in liquid culture using miRNeasy Micro Kit (QIAGEN), and DNase treatment was performed using RNase-free DNase Set (QIAGEN), according to the manufacturer’s instructions. Total RNA was used for library preparation with NEBNext Ultra-low input II Directional RNA Library Prep Kit (Illumina) and sequenced on an Illumina HiSeq with 2×150 bp sequencing configuration (Illumina).

### RNA-seq analysis

High quality reads, obtained after reads quality inspection and adapter trimming with Trim Galore (https://www.bioinformatics.babraham.ac.uk/projects/trim_galore/) were aligned to the human reference genome (GENCODE GRCh38 primary assembly) with STAR release 2.7.6a. Gene expression quantification was obtained with featureCounts v2.0.1, using the GENCODE annotation (version v35 primary assembly). Raw counts were normalized by library size, and the differential expression (DE) analysis was performed with the DESeq2 package, testing for the contrast OLD vs. YOUNG. Finally, Benjamini-Hochberg adjusted *P* values were used to retain significant differentially expressed genes (FDR < 0.05). For the Gene Set Enrichment Analysis (GSEA)^94^ genes were ranked by Log-Fold Change and analyzed with the *gsePathway* function from clusterProfiler, considering the Reactome database, retaining gene categories with FDR < 0.05. All figures were plotted with ggplot2 in R.

### Colony forming unit assay

CFU assay was performed at the indicated time points plating 800 cells in methylcellulose-based medium (MethoCult H4434, StemCell Technologies) supplemented with penicillin and streptomycin. Two weeks after plating, colonies were counted in blind and identified as myeloid and erythroid colonies according to morphological criteria.

### *In vitro* growth curve of young and old CD34+ cells

Human CD34+ cells were seeded at the concentration of 5*10^5^ CD34+ cells/ml in serum-free StemSpan medium (StemCell Technologies) supplemented with penicillin, streptomycin, glutamine and human early-acting cytokines (SCF 300 ng/ml, Flt3-L 300 ng/ml, TPO 100 ng/ml, and IL-3 60 ng/ml; all purchased from Peprotech). All cultures were kept at 37°C in a 5% CO2 water jacket incubator (Thermo Scientific) and cell count was assessed every 24h hours.

### Ki67 staining and flow cytometry analysis

Cells were washed and fixed using BD Cytofix buffer (Cat. #554655), washed and permeabilized with BD Perm 2 (Cat. # 347692), washed and stained with FITC-conjugated Ki67 antibody (BD Biosciences, 556026). The cells were then analysed on a BD LSR-Fortessa cytometer (BD Biosciences).

### Cell Trace Proliferation Assay

Bulk HSPCs or FACS-sorted HSPC subsets from young and aged healthy donors were washed and incubated with CellTrace Violet (ThermoFisher, C34557) 5 mM for 20 minutes at 37°, protected from light. Cell sorting was performed using FACS Aria Fusion (BD Biosciences). After incubation, cells were washed and resuspended in a fresh pre-warmed complete culture medium and cultured in serum-free StemSpan medium (StemCell Technologies) supplemented with penicillin, streptomycin, glutamine and hSCF 300 ng/ml, hFlt3-L 300 ng/ml, hTPO 100 ng/ml, and hIL-3 60 ng/ml (all purchased from Peprotech). Cells were collected every 24h to measure the cell divisions through flow cytometry. Cells were acquired to BD LSRFortessa Cytometer (BD Biosciences).

### Gene expression analysis

For gene expression analyses, total RNA was extracted using either miRNeasy Micro Kit (QIAGEN) or RNeasy Plus Micro Kit (QIAGEN), according to the manufacturer’s instructions and DNase treatment was performed using RNase-free DNase Set (QIAGEN). cDNA was synthetized with iScript cDNA Synthesis Kit (Bio-Rad). For selected analyses, cDNA was then pre-amplified using TaqMan PreAmp Master Mix (ThermoFisher) and used for q-PCR in a Viia7 Real-time PCR thermal cycler using both Fast SYBR Green Master Mix (ThermoFisher), after standard curve method optimization to reach the 100% primer efficiency for each couple of primers listed in Table S2. The relative expression of each target gene was first normalized to *GUSB* or β*2M* housekeeping genes expression and then represented as 2320 cl:838-ΔCt relative to the indicated control conditions.

### Immunofluorescence analysis

Multi-test slides (10 well, MP Biomedicals) were treated for 20’ with Poly-L-lysine solution (Sigma-Aldrich) at 1mg/ml concentration. After two washes with DPBS solution, approximately 0.5/1×10^5^ cells were seeded on covers for 20’ and fixed with 4% paraformaldehyde (Santa Cruz Biotechnology) for other 20’. Cells were then permeabilized with 0.2% Triton X100. After blocking with 0.5% BSA and 0.2% fish gelatin in DPBS, cells were probed with the indicated primary antibodies. After primary antibodies incubation (Anti-phospho RPA (Ser33) Antibody, Merck) cells were washed three times with DPBS and incubate with Alexa 488-, 568- and/or 647-labeled secondary antibodies (Invitrogen). Nuclear DNA was stained with DAPI (Sigma-Aldrich) and covers were mounted with Aqua-Poly/Mount solution (Polysciences. Inc.) on glass slides (Bio-Optica). Fluorescent images were acquired using Leica SP2 and Leica SP5 Confocal microscopes. Where indicated, quantification of DNA damage response nuclear foci in immunofluorescence images was conducted using ImageJ64 (version 1.47).

### Transplantation experiments of activated human HSPC across aging and for the *in vivo* model of proliferation stress

Mouse studies were conducted according to protocols approved by the San Raffaele Scientific Institute and the Italian Ministry of Health (IACUC, #872). NOD.Cg-Prkdc^scid^ IL2rγ^tm1Wjl^/SzJ (NSG) mice were purchased from The Jackson Laboratory (cat. #005557 Jackson Laboratories, Bar Harbor, ME). All animals were maintained in the SPF animal facility at IRCCS Ospedale San Raffaele. The handling of the animals was performed by trained personnel, with the aim of minimizing the degree of stress and suffering and the period of constraint.

For transplantation, 2.5×10^5^ BM-CD34+ cells isolated from young and old healthy donors or different doses of CB-CD34+ (50×10^3^; 10×10^3^; 3×10^3^) were injected intravenously into 8-weeks old female sub-lethally irradiated NSG mice (150-180 cGy). Mice were monitored three times a week by weight assessment and general appearance, taking into consideration motility, fur conditions, kyphosis, and other signs of disease. Human CD45+ cell engraftment and blood lineages reconstitution were monitored by serial collection of blood from the mouse tail and at time of sacrifice BM, spleen and distal organs were harvested. Human cell engraftment in murine organs was assessed by applying WBD protocol (see “Whole Blood Dissection Staining” section). Before WBD procedure, cells were incubated with a mouse FcR blocking reagent (BD Biosciences, dilution 1:100).

### BrdU and EdU *in vivo* administration and staining procedure

For cell proliferation analysis, EdU and BrdU double labelling was used as previously described^66^. Both nucleotide analogues are incorporated during DNA synthesis and, when combined, allow to analyse cell cycle kinetics. Pulse-chase timings were previously optimized^66^ to achieve consistent *in vivo* incorporation of EdU and BrdU, and later detection in the cell population of interest. EdU (1 mg/mouse, Component A, Click-iT EdU Alexa Fluor 647, ThermoFisher, C10635) and BrdU (2 mg/mouse, Sigma-Aldrich, B5002-100MG) were subsequently administrated via tail vein injections (2h apart); 30 min after BrDU injection, mice were culled, and BM cells were isolated to perform human cell staining. The staining procedure was performed as follows:

- Surface staining of BM cells with labeled with the following fluorescent antibodies:
- Mouse anti-human CD45-BUV395 (Clone: HI30; BD Biosciences, 563792), Verified Reactivity: Human (QC Testing); Application: Flow cytometry (Routinely Tested). Dilution: 1:33
- Mouse anti-human CD3-BV605 (Clone: OKT3; Biolegend, 317322), Verified Reactivity: Human; Application: Flow cytometric analysis of antibody surface-stained cells. Dilution: 1:50
- Mouse anti-human CD56-PC5 (Clone: 5.1H11; Biolegend, 362516), Verified Reactivity: Human; Application: Flow cytometric analysis of antibody surface-stained cells. Dilution: 1:50
- Mouse anti-human CD14-PE (Clone: M5E2; Biolegend, 301806), Verified Reactivity: Human, Cynomolgus, Rhesus; Application: Flow cytometric analysis of antibody surface-stained cells. Dilution: 1:50
- Mouse anti-human CD34-BV421 (Clone: 561; Biolegend, 343610), Verified Reactivity: Human; Application: Flow cytometric analysis of antibody surface-stained cells. Dilution: 1:20
- Mouse anti-human CD15-APCfire750 (Clone: W6D3; Biolegend, 323041), Verified Reactivity: Human; Application: Flow cytometric analysis of antibody surface-stained cells. 1:50
- Mouse anti-human CD19-APCR700 (Clone: SJ25C1; BD Biosciences, 659121), Verified Reactivity: Human; Application: Flow cytometry. Dilution: 1:20
- Mouse anti-human CD38-BV510 (Clone: HB-7; Biolegend, 356612), Verified Reactivity: Human (QC Testing); Application: Flow cytometry (Routinely Tested). Dilution: 1:40
- Mouse anti-human CD11c-BV650 (Clone: B-ly6; BD Biosciences, 563404), Verified Reactivity: Human (QC Testing); Application: Flow cytometry (Routinely Tested). Dilution: 1:20
- Fixation and permeabilization followed by EdU staining (Click-iT EdU Alexa Fluor 647, ThermoFisher, C10635) according to manufactures’ instructions.
- Permeabilization using Cytoperm Buffer PLUS (BD Biosciences) 10 mins on ice followed by permeabilization through CytoFix/CytoPerm kit solution (BD Biosciences) 5 mins at room temperature
- DNAse treatment (Qiagen) for 1 hour at 37°C
- Staining with anti-BrDU-Alexa488 antibody, Clone: 3D4, Biolegend, 364105, Application: Intracellular Flow cytometry (Routinely Tested). Dilution: 1:50.

Samples were acquired at BD LSR-Fortessa (BD Biosciences) at 400 event/sec rate.

### Neutral and Alkaline Comet Assays

Purified HSPCs from CD34+ cells isolated from mice transplanted with distinct doses of CD34+ cells at 20 weeks were suspended at 1 x 10^5^ cells/ml in ice cold 1X DPBS and mixed with molten Comet LMAgarose (Trevigen, MD) at a ratio of 1:10 (v/v) and immediately pipetted onto CometSlides (Trevigen, MD) and placed at +4°C. Once solidified, the slides were immersed in pre-chilled Lysis Solution (Trevigen, MD) overnight at +4°C. For *Neutral comet assay,* after lysis, slides were immersed in Neutral Electrophoresis Buffer pH 8 (50 mM Tris, 150 mM Sodium Acetate) for 1 hour at +4°C and then electrophoresed in Neutral Electrophoresis Buffer pH 8 (100 mM Tris, 300 mM Sodium Acetate) at 300 mA for 40 min. Slides were gently immersed in DNA Precipitation Solution (375 mM Ammonium Acetate in Ethanol 95%) for 30 minutes at RT and fixed in 70% ethanol for 30 min. For *Alkaline comet assay*, after lysis, slides were immersed in freshly prepared Alkaline Unwinding Solution pH > 13 (300 mM NaOH, 1mM EDTA) for 1 hour at +4°C and then electrophoresed in Alkaline Electrophoresis Solution pH > 13 (300 mM NaOH, 1mM EDTA) at 300 mA for 20 min. Slides were washed twice in ddH2O and fixed in 70% ethanol for 5 min. Comets were stained with SYBR Safe (Invitrogen) for 30 min at RT. All steps were conducted in the dark to prevent additional DNA damage. Comets were analysed using a Nikon Eclipse E600 microscope and a Nikon-DS-RI2 camera. At least 80 nuclei for each individual donor were analysed with CASP software to determine “Olive Tail Moments” of individual nuclei.

### Statistical analyses

Statistical test and *p* values were specified in each figure legend or in within figure graphs. To assess the correlation between two variables the Spearman index was calculated and a regression line(with 95% confidence intervals) was estimated after assessing the normality of residuals as appropriate. Non-parametric tests were used for comparing two or more groups when continuous variables were considered. The analyses were performed using Prism v9.1.0 software (GraphPad). For the longitudinal analyses of growth curve of cultured cells, a linear mixed-effects (LME) model has been applied. To better capture growth dynamics, group indicator (young *vs.* old) and time variable (a categorical variable taking 5 values, from 0 to 96 hours) were entered in the model as main effects as well as in interaction. To account for donor-specific heterogeneity, a random effect was specified on donors’ ID, thus leading to estimate random intercept models. This modelling approach properly accounts for the dependency among observations arising from measuring the same unit more than once and over time. Logarithmic transformation of the outcome variable was considered to satisfy underlying model assumptions. Analyses were performed using R statistical software (version 4.0.4). Significance level was set at 0.05. LME were estimated using the *nlme* package and post-hoc comparisons, to compare groups at a fixed time point, have been performed using *emmeans* package.

## Acknowledgments

This work was supported by Fondazione Telethon (TIGET Core Grant B2, A.A TGT21016, A.A and S.S.), the Italian Ministero della Salute (GR-2019-12369499, S.S.) and Else Kröner-Fresenius-Stiftung (EKFS) prize (A.A.). Research in Di Micco’s lab is supported by a My First AIRC Grant MFAG 2019–PI ID.23321, by Telethon (TIGET Core grant E5 and TGT21018), by a Career Development Award from Human Frontier Science Program, by the Interstellar Initiative on Healthy Longevity from the New York Academy of Sciences and the Japan Agency for Medical Research and Development, by the ASH Global Research Award, by the New York Stem Cell Foundation,the European Research Council (Consolidator grant 101003186, ReviveSTEM), and the X-PAND Pathfinder European Commission grant (G.A. 101070950). A.C. is an early career scientist in the European Hematology Association and the American Society of Hematology Translational Research Training in Hematology (EHA and ASH TRTH). T.T. is enrolled in the EHA-EMBL/EBI computational biology training in hematology. S.Z.X. is supported by funds from the Princess Margaret Cancer Foundation. We thank F. Ciceri, ME Bernardo and all medical and nursing staff at Raffaele Scientific Institute and the San Raffaele Stem Cell Programme; we thank C. Villa and S. Di Terlizzi of the Flow Cytometry Resource, Advanced Cytometry Technical Applications Laboratory (FRACTAL) at Ospedale San Raffaele for cell sorting and technical help with instrumentation; we thank C. Brombin of University Center for Statistics in the Biomedical Science (CUSSB) for LME statistical models. E.L. L.d.V and P.Q. conducted this study as partial fulfilment of their Ph.D. in Molecular Medicine, Cellular and Molecular Physiopathology Program, San Raffaele University, Milan, Italy. R.D.M. is a New York Stem Cell Foundation Robertson Investigator.

## Authorship contributions

E.L. and S.S. performed phenotypic characterization, *in vitro* and *in vivo* assays, collected and interpreted the data, and wrote the manuscript; L.B.-R. performed phenotypic characterization and isolation of HSPC subpopulations, performed *in vitro* and *in vivo* assays and analysed the data; L.G.P., K.B.K., and S.Z.X performed phenotypic characterization and isolation of HSPC subpopulations, quantitative xenotransplantation, and collected data. L.G.P. performed ATAC-seq sample preparation; A.M. performed analyses of the ATAC-seq dataset; S.Z.X. analysed data; P.Q K.G. and R.J.H. performed *in vivo* assays; T.T. performed analyses of RNAseq dataset; L.d.V. performed immunofluorescence analyses on *in vivo* experiments; E.L.F. performed Comet assay and analyses; A.C. and G.F. generated RNAseq dataset; P.C, M.O., A.G.A.N., E.F.F. provided human BM samples from aged donors; S.B and I.M. supervised RNAseq analyses; S.Z.X., A.A. and R.D.M interpreted data, supervised team members and contributed to writing-review and editing of the manuscript. R.D.M conceived the study and secured funding.

## Data and Code availability

All the data generated in this study are in the process of being submitted to NCBI/GEO database. The code used to process and to generate the images of RNAseq data in this manuscript is available at: http://www.bioinfotiget.it/gitlab/custom/LetteraScala_Ageing/bulk_RNAseq.

## REFERENCES

1. Sudo, K., Ema, H., Morita, Y. & Nakauchi, H. Age-associated characteristics of murine hematopoietic stem cells [In Process Citation]. J Exp Med 192, (2000).

2. de Haan, G. & Lazare, S. S. Aging of hematopoietic stem cells. Blood 131, 479–487 (2018).

3. Morrison, S. J., Wandycz, A. M., Akashi, K., Globerson, A. & Weissman, I. L. The aging of hematopoietic stem cells. Nat Med 2, 1011–6 (1996).

4. Young, K. et al. Decline in IGF1 in the bone marrow microenvironment initiates hematopoietic stem cell aging. Cell Stem Cell 28, 1473–1482.e7 (2021).

5. Oh, J., Lee, Y. D. & Wagers, A. J. Stem cell aging: mechanisms, regulators and therapeutic opportunities. Nat Med 20, 870–80 (2014).

6. Sun, D. et al. Epigenomic profiling of young and aged HSCs reveals concerted changes during aging that reinforce self-renewal. Cell Stem Cell 14, 673–88 (2014).

7. Rossi, D. J. et al. Cell intrinsic alterations underlie hematopoietic stem cell aging. Proc Natl Acad Sci U S A 102, 9194–9 (2005).

8. Beerman, I., Seita, J., Inlay, M. A., Weissman, I. L. & Rossi, D. J. Quiescent hematopoietic stem cells accumulate DNA damage during aging that is repaired upon entry into cell cycle. Cell Stem Cell 15, 37–50 (2014).

9. Luchsinger, L. L., de Almeida, M. J., Corrigan, D. J., Mumau, M. & Snoeck, H.-W. Mitofusin 2 maintains haematopoietic stem cells with extensive lymphoid potential. Nature 529, 528–31 (2016).

10. Simsek, T. et al. The distinct metabolic profile of hematopoietic stem cells reflects their location in a hypoxic niche. Cell Stem Cell 7, 380–90 (2010).

11. Morita, Y., Ema, H. & Nakauchi, H. Heterogeneity and hierarchy within the most primitive hematopoietic stem cell compartment. J Exp Med 207, 1173–82 (2010).

12. Bernitz, J. M., Kim, H. S., MacArthur, B., Sieburg, H. & Moore, K. Hematopoietic Stem Cells Count and Remember Self-Renewal Divisions. Cell 167, 1296–1309.e10 (2016).

13. Kirschner, K. et al. Proliferation Drives Aging-Related Functional Decline in a Subpopulation of the Hematopoietic Stem Cell Compartment. Cell Rep 19, 1503–1511 (2017).

14. Walter, D. et al. Exit from dormancy provokes DNA-damage-induced attrition in haematopoietic stem cells. Nature 520, 549–552 (2015).

15. Beerman, I. et al. Proliferation-dependent alterations of the DNA methylation landscape underlie hematopoietic stem cell aging. Cell Stem Cell 12, 413–25 (2013).

16. Flach, J. et al. Replication stress is a potent driver of functional decline in ageing haematopoietic stem cells. Nature 512, 198–202 (2014).

17. Wang, M. et al. Genotoxic aldehyde stress prematurely ages hematopoietic stem cells in a p53-driven manner. Mol Cell 83, 2417–2433.e7 (2023).

18. Liang, Y., Van Zant, G. & Szilvassy, S. J. Effects of aging on the homing and engraftment of murine hematopoietic stem and progenitor cells. Blood 106, 1479–87 (2005).

19. Chung, S. S. & Park, C. Y. Aging, hematopoiesis, and the myelodysplastic syndromes. Blood Adv 1, 2572–2578 (2017).

20. van Galen, P. et al. Reduced lymphoid lineage priming promotes human hematopoietic stem cell expansion. Cell Stem Cell 14, 94–106 (2014).

21. Pang, W. W., Schrier, S. L. & Weissman, I. L. Age-associated changes in human hematopoietic stem cells. Semin Hematol 54, 39–42 (2017).

22. von Bonin, M. et al. Clonal hematopoiesis and its emerging effects on cellular therapies. Leukemia 35, 2752–2758 (2021).

23. Jaiswal, S. & Ebert, B. L. Clonal hematopoiesis in human aging and disease. Science (1979) 366, (2019).

24. Signer, R. A. J., Montecino-Rodriguez, E., Witte, O. N., McLaughlin, J. & Dorshkind, K. Age-related defects in B lymphopoiesis underlie the myeloid dominance of adult leukemia. Blood 110, 1831–1839 (2007).

25. Vas, V., Wandhoff, C., Dörr, K., Niebel, A. & Geiger, H. Contribution of an Aged Microenvironment to Aging-Associated Myeloproliferative Disease. PLoS One 7, e31523 (2012).

26. Henry, C. J., Marusyk, A., Zaberezhnyy, V., Adane, B. & DeGregori, J. Declining lymphoid progenitor fitness promotes aging-associated leukemogenesis. Proceedings of the National Academy of Sciences 107, 21713–21718 (2010).

27. Stauder, R., Valent, P. & Theurl, I. Anemia at older age: etiologies, clinical implications, and management. Blood 131, 505–514 (2018).

28. Shlush, L. I. Age-related clonal hematopoiesis. Blood 131, 496–504 (2018).

29. Avagyan, S. & Zon, L. I. Clonal hematopoiesis and inflammation – the perpetual cycle. Trends Cell Biol 33, 695–707 (2023).

30. Belluschi, S. et al. Myelo-lymphoid lineage restriction occurs in the human haematopoietic stem cell compartment before lymphoid-primed multipotent progenitors. Nat Commun 9, 4100 (2018).

31. Zhang, Y. W. et al. Hyaluronic acid-GPRC5C signalling promotes dormancy in haematopoietic stem cells. Nat Cell Biol 24, 1038–1048 (2022).

32. García-Prat, L. et al. TFEB-mediated endolysosomal activity controls human hematopoietic stem cell fate. Cell Stem Cell 28, 1838–1850.e10 (2021).

33. Milyavsky, M. et al. A distinctive DNA damage response in human hematopoietic stem cells reveals an apoptosis-independent role for p53 in self-renewal. Cell Stem Cell 7, 186–97 (2010).

34. Laurenti, E. & Göttgens, B. From haematopoietic stem cells to complex differentiation landscapes. Nature 553, 418–426 (2018).

35. Takayama, N. et al. The Transition from Quiescent to Activated States in Human Hematopoietic Stem Cells Is Governed by Dynamic 3D Genome Reorganization. Cell Stem Cell 28, 488–501.e10 (2021).

36. Kaufmann, K. B. et al. A latent subset of human hematopoietic stem cells resists regenerative stress to preserve stemness. Nat Immunol 22, 723–734 (2021).

37. Xie, S. Z. et al. Sphingosine-1-phosphate receptor 3 potentiates inflammatory programs in normal and leukemia stem cells to promote differentiation. Blood Cancer Discov 2, 32–53 (2021).

38. Notta, F. et al. Distinct routes of lineage development reshape the human blood hierarchy across ontogeny. Science 351, aab2116 (2016).

39. Notta, F. et al. Isolation of single human hematopoietic stem cells capable of long-term multilineage engraftment. Science 333, 218–21 (2011).

40. Laurenti, E. et al. CDK6 levels regulate quiescence exit in human hematopoietic stem cells. Cell Stem Cell 16, 302–13 (2015).

41. Hammond, C. A., Wu, S. W., Wang, F., MacAldaz, M. E. & Eaves, C. J. Aging alters the cell cycle control and mitogenic signaling responses of human hematopoietic stem cells. Blood 141, 1990–2002 (2023).

42. Amoah, A. et al. Aging of human hematopoietic stem cells is linked to changes in Cdc42 activity. Haematologica 107, 393–402 (2022).

43. Jiang, P. et al. Immune reconstitution and survival of patients after allogeneic hematopoietic stem cell transplantation from older donors. Clin Transplant 37, (2023).

44. Duda, K. et al. Allogeneic hematopoietic stem cell transplantation remains a feasible approach for elderly with acute myeloid leukemia: a 10-year experience. Ann Hematol 102, 1907–1914 (2023).

45. Mehta, R. S. et al. Haploidentical vs matched unrelated donors for patients with ALL: donor age matters more than donor type. Blood Adv 7, 1594–1603 (2023).

46. Artz, A. S. From Biology to Clinical Practice: Aging and Hematopoietic Cell Transplantation. Biology of Blood and Marrow Transplantation 18, S40–S45 (2012).

47. Kollman, C. et al. Donor characteristics as risk factors in recipients after transplantation of bone marrow from unrelated donors: the effect of donor age. Blood 98, 2043–2051 (2001).

48. Basso-Ricci, L., et al. Multiparametric Whole Blood Dissection: A one-shot comprehensive picture of the human hematopoietic system. Cytometry Part A 91, 952–965 (2017).

49. Pang, W. W. et al. Human bone marrow hematopoietic stem cells are increased in frequency and myeloid-biased with age. Proc Natl Acad Sci U S A 108, 20012–7 (2011).

50. Rundberg Nilsson, A., Soneji, S., Adolfsson, S., Bryder, D. & Pronk, C. J. Human and Murine Hematopoietic Stem Cell Aging Is Associated with Functional Impairments and Intrinsic Megakaryocytic/Erythroid Bias. PLoS One 11, e0158369 (2016).

51. Zhang, L., Mack, R., Breslin, P. & Zhang, J. Molecular and cellular mechanisms of aging in hematopoietic stem cells and their niches. J Hematol Oncol 13, 157 (2020).

52. Kuranda, K. et al. Age-related changes in human hematopoietic stem/progenitor cells. Aging Cell 10, 542–546 (2011).

53. Wang, J. C., Doedens, M. & Dick, J. E. Primitive human hematopoietic cells are enriched in cord blood compared with adult bone marrow or mobilized peripheral blood as measured by the quantitative in vivo SCID-repopulating cell assay. Blood 89, 3919–24 (1997).

54. Xu, C.-X. et al. ETV2/ER71 regulates hematopoietic regeneration by promoting hematopoietic stem cell proliferation. J Exp Med 214, 1643–1653 (2017).

55. Pimanda, J. E. et al. Gata2, Fli1, and Scl form a recursively wired gene-regulatory circuit during early hematopoietic development. Proc Natl Acad Sci U S A 104, 17692–7 (2007).

56. Kruse, E. A. et al. Dual requirement for the ETS transcription factors Fli-1 and Erg in hematopoietic stem cells and the megakaryocyte lineage. Proc Natl Acad Sci U S A 106, 13814–9 (2009).

57. Iaquinta, P. J. & Lees, J. A. Life and death decisions by the E2F transcription factors.

58. Luc, S. et al. Bcl11a Deficiency Leads to Hematopoietic Stem Cell Defects with an Aging-like Phenotype. Cell Rep 16, 3181–3194 (2016).

59. Amsellem, S. et al. Ex vivo expansion of human hematopoietic stem cells by direct delivery of the HOXB4 homeoprotein. Nat Med 9, 1423–1427 (2003).

60. Sauvageau, G. et al. Overexpression of HOXB4 in hematopoietic cells causes the selective expansion of more primitive populations in vitro and in vivo. Genes Dev 9, 1753–65 (1995).

61. Miller, C. L. & Eaves, C. J. Expansion in vitro of adult murine hematopoietic stem cells with transplantable lympho-myeloid reconstituting ability. Proc Natl Acad Sci U S A 94, 13648– 53 (1997).

62. Conneally, E., Cashman, J., Petzer, A. & Eaves, C. Expansion *in vitro* of transplantable human cord blood stem cells demonstrated using a quantitative assay of their lympho-myeloid repopulating activity in nonobese diabetic– *scid/scid* mice. Proceedings of the National Academy of Sciences 94, 9836–9841 (1997).

63. Bhatia, M. et al. Quantitative analysis reveals expansion of human hematopoietic repopulating cells after short-term ex vivo culture. J Exp Med 186, 619–24 (1997).

64. Ferrari, G., Thrasher, A. J. & Aiuti, A. Gene therapy using haematopoietic stem and progenitor cells. Nat Rev Genet 22, 216–234 (2021).

65. Ferrari, S. et al. Genetic engineering meets hematopoietic stem cell biology for next-generation gene therapy. Cell Stem Cell 30, 549–570 (2023).

66. Akinduro, O. et al. Proliferation dynamics of acute myeloid leukaemia and haematopoietic progenitors competing for bone marrow space. Nat Commun 9, 519 (2018).

67. Olive, P. L. & Banáth, J. P. The comet assay: a method to measure DNA damage in individual cells. Nat Protoc 1, 23–29 (2006).

68. Ciccia, A. & Elledge, S. J. The DNA damage response: making it safe to play with knives. Mol Cell 40, 179–204 (2010).

69. Kovtonyuk, L. V., Fritsch, K., Feng, X., Manz, M. G. & Takizawa, H. Inflamm-Aging of Hematopoiesis, Hematopoietic Stem Cells, and the Bone Marrow Microenvironment. Front Immunol 7, (2016).

70. Franceschi, C. et al. Inflammaging and anti-inflammaging: A systemic perspective on aging and longevity emerged from studies in humans. Mech Ageing Dev 128, 92–105 (2007).

71. Gowanlock, Z., Sriram, S., Martin, A., Xenocostas, A. & Lazo-Langner, A. Erythropoiesis-stimulating agents in elderly patients with anemia: response and cardiovascular outcomes. Blood Adv 1, 1538–1545 (2017).

72. Zeng, A. G. X. et al. A hematopoietic stem cell subset that retains memory of prior inflammatory stress accumulates in aging and clonal hematopoiesis. bioRxiv 2023.09.11.557271 (2023) doi:10.1101/2023.09.11.557271.

73. McEwen, B. S. Interacting mediators of allostasis and allostatic load: towards an understanding of resilience in aging. Metabolism 52, 10–16 (2003).

74. Cirovic, B. et al. BCG Vaccination in Humans Elicits Trained Immunity via the Hematopoietic Progenitor Compartment. Cell Host Microbe 28, 322–334.e5 (2020).

75. de Laval, B. et al. C/EBPβ-Dependent Epigenetic Memory Induces Trained Immunity in Hematopoietic Stem Cells. Cell Stem Cell 26, 657–674.e8 (2020).

76. Kaufmann, E. et al. BCG Educates Hematopoietic Stem Cells to Generate Protective Innate Immunity against Tuberculosis. Cell 172, 176–190.e19 (2018).

77. Naik, S. & Fuchs, E. Inflammatory memory and tissue adaptation in sickness and in health. Nature 607, 249–255 (2022).

78. Bogeska, R. et al. Inflammatory exposure drives long-lived impairment of hematopoietic stem cell self-renewal activity and accelerated aging. Cell Stem Cell 29, 1273–1284.e8 (2022).

79. Naik, S. et al. Inflammatory memory sensitizes skin epithelial stem cells to tissue damage. Nature 550, 475–480 (2017).

80. Jacobs, K. et al. Stress-triggered hematopoietic stem cell proliferation relies on PrimPol-mediated repriming. Mol Cell 82, 4176–4188.e8 (2022).

81. Kruta, M. et al. Hsf1 promotes hematopoietic stem cell fitness and proteostasis in response to ex vivo culture stress and aging. Cell Stem Cell 28, 1950–1965.e6 (2021).

82. Keyvani Chahi, A., et al. PLAG1 dampens protein synthesis to promote human hematopoietic stem cell self-renewal. Blood 140, 992–1008 (2022).

83. Papa, L. et al. Ex vivo human HSC expansion requires coordination of cellular reprogramming with mitochondrial remodeling and p53 activation. Blood Adv 2, 2766–2779 (2018).

84. Xie, S. Z. et al. Sphingolipid Modulation Activates Proteostasis Programs to Govern Human Hematopoietic Stem Cell Self-Renewal. Cell Stem Cell 25, 639–653.e7 (2019).

85. Lahav, M. et al. Nonmyeloablative Conditioning Does Not Prevent Telomere Shortening after Allogeneic Stem Cell Transplantation. Transplantation 80, 969–976 (2005).

86. Perna, S. et al. Performance of Edmonton Frail Scale on frailty assessment: its association with multi-dimensional geriatric conditions assessed with specific screening tools. BMC Geriatr 17, 2 (2017).

87. Aguilar-Navarro, A. G. et al. Human Aging Alters the Spatial Organization between CD34+ Hematopoietic Cells and Adipocytes in Bone Marrow. Stem Cell Reports 15, 317–325 (2020).

88. Hu, Y. & Smyth, G. K. ELDA: extreme limiting dilution analysis for comparing depleted and enriched populations in stem cell and other assays. J Immunol Methods 347, 70–8 (2009).

89. Buenrostro, J. D., Giresi, P. G., Zaba, L. C., Chang, H. Y. & Greenleaf, W. J. Transposition of native chromatin for fast and sensitive epigenomic profiling of open chromatin, DNA-binding proteins and nucleosome position. Nat Methods 10, 1213–8 (2013).

90. Li, H. & Durbin, R. Fast and accurate short read alignment with Burrows-Wheeler transform. Bioinformatics 25, 1754–60 (2009).

91. Zhang, Y. et al. Model-based Analysis of ChIP-Seq (MACS). Genome Biol 9, R137 (2008).

92. Gaujoux, R. & Seoighe, C. A flexible R package for nonnegative matrix factorization. BMC Bioinformatics 11, 367 (2010).

93. Heinz, S. et al. Simple Combinations of Lineage-Determining Transcription Factors Prime cis-Regulatory Elements Required for Macrophage and B Cell Identities. Mol Cell 38, 576– 589 (2010).

94. Subramanian, A. et al. Gene set enrichment analysis: A knowledge-based approach for interpreting genome-wide expression profiles. Proceedings of the National Academy of Sciences 102, 15545–15550 (2005).

