## Supplementary DATA file for "Molecular and phenotypic blueprint of the hematopoietic compartment reveals proliferation stress as a driver of age-associated human stem cell dysfunctions"

#### Supplementary Data items

This file contains:

- 7 Supplementary Figures + Legends

#### Supplementary Figure 1

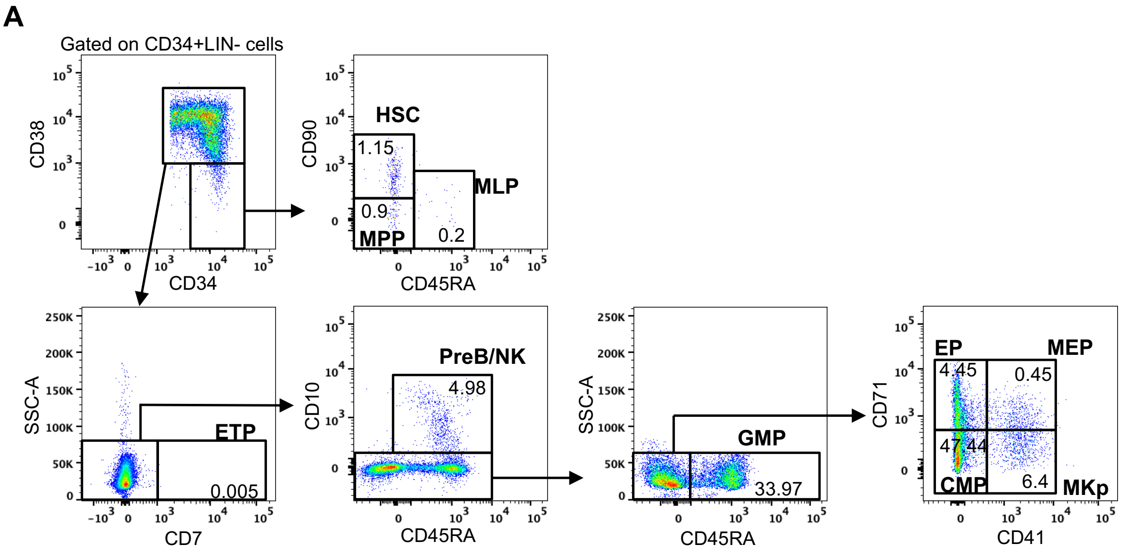


**Supplementary Figure 1. Representative plots and gating strategy for human HSPC subpopulations.** Representative plots of CD34+LIN- composition and gating strategy used for the identification of HSPC subpopulations from a representative young donor. Numbers indicate the frequencies of each HSPC subset on CD34+LIN- cells.

### Supplementary Figure 2

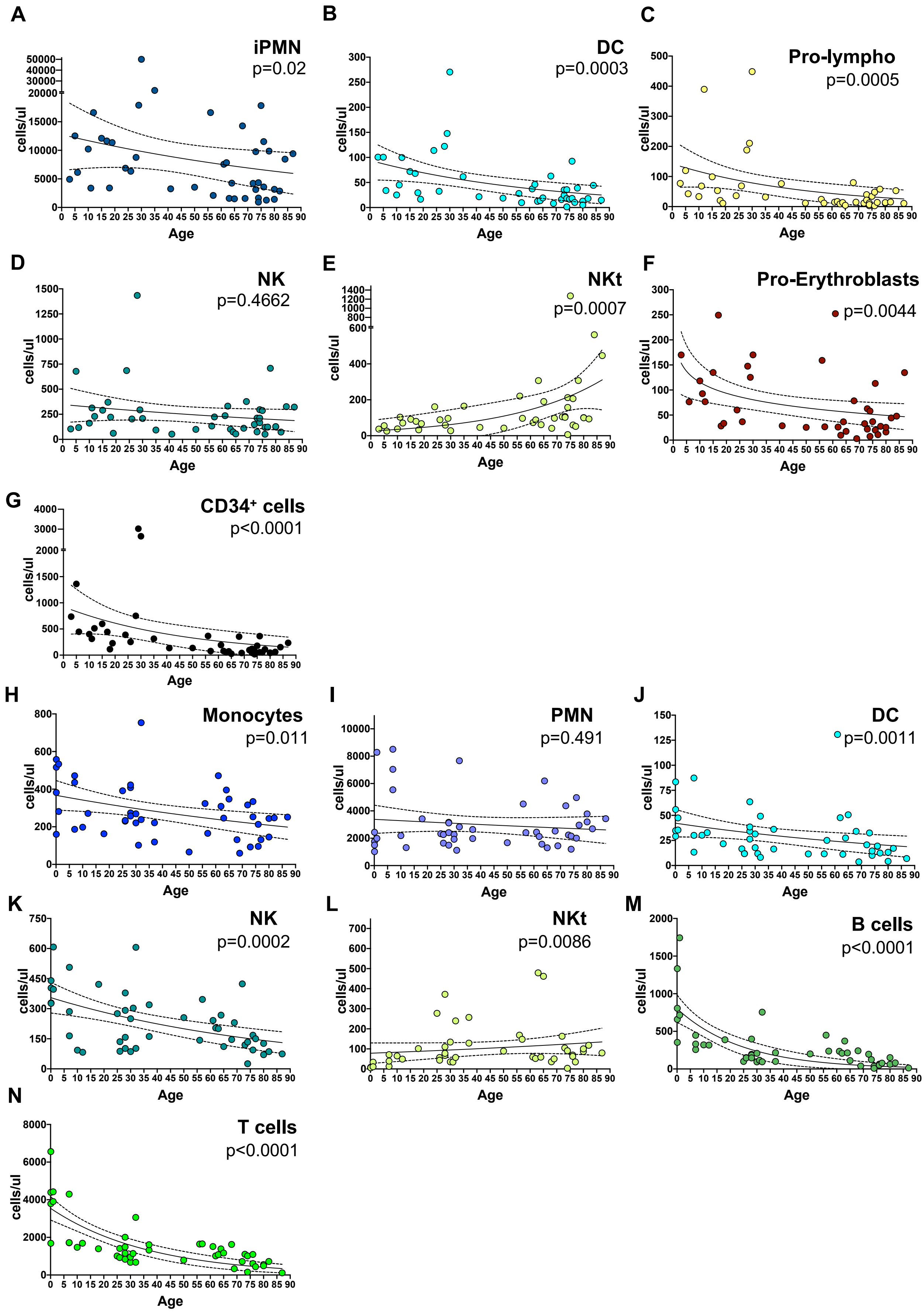

**Supplementary Figure 2. A-G.** Correlation between the absolute count of each cell subtype (expressed as cells/µL) in the BM and the age of the 45 subjects analyzed. **H-N.** Correlation between the absolute count of each cell subtype (expressed as cells/µL) in the PB and the age of the 56 subjects analyzed. (iPMN, immature Polymorphonucleated cells; PMN, polymorphonucleated cells; DCs, dendritic cells; Pro-lympho, lymphocytes progenitors; NK, natural killer; NKt, natural killer T cells; Pro-Erythroblasts, erythroblast progenitors).

### Supplementary Figure 3

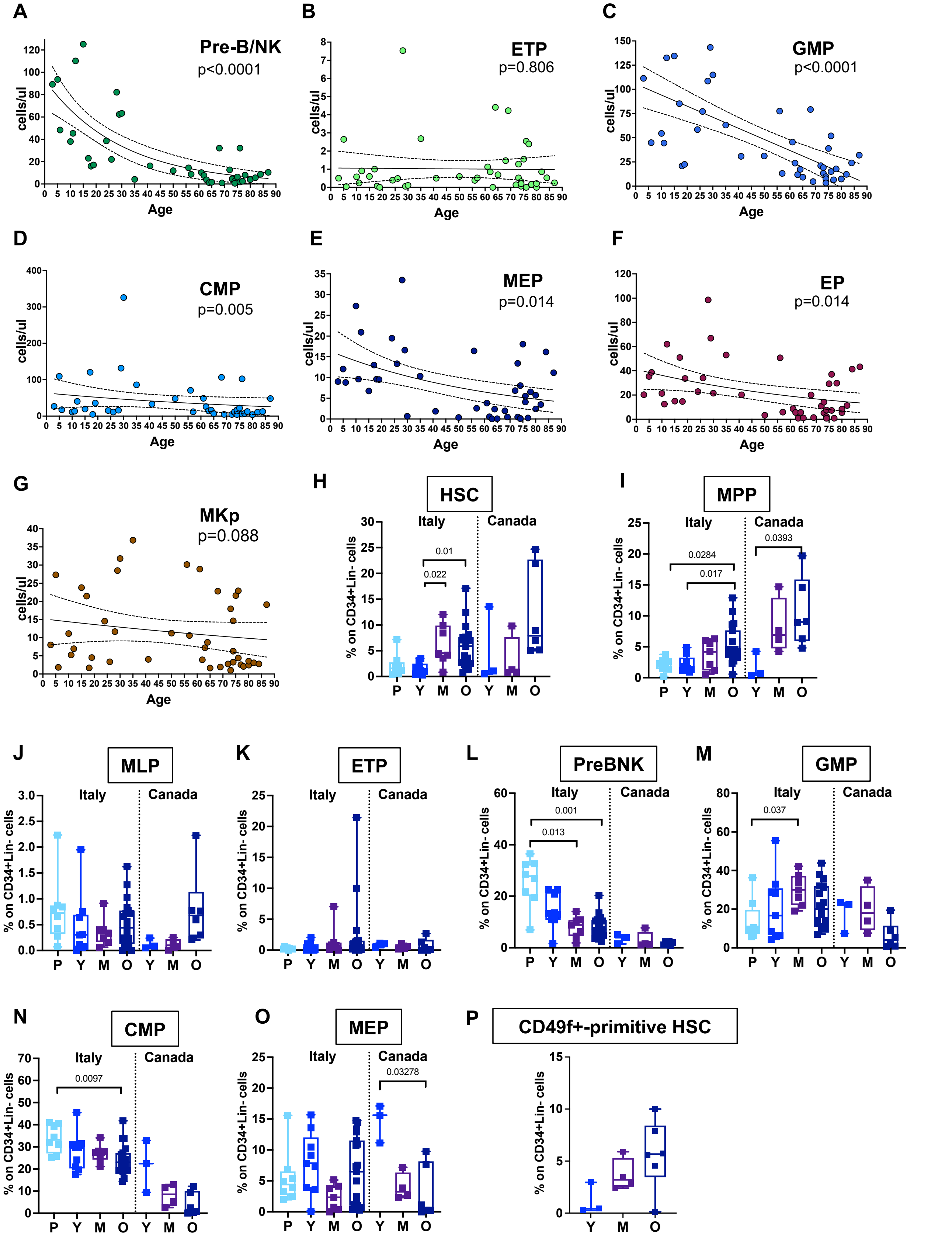


**Supplementary Figure 3. A-G.** Correlation between the absolute count of each HSPC subtype (expressed as cells/μL) in the BM and the age of the 45 subjects analyzed. **H-O.** Frequencies of distinct HSPC subpopulations within CD34+LIN- compartment in healthy donors from different ages in the Italian cohort (P=Pediatric <18 years old, n=8; Y= young 18-35 years old, n=10; M=Middle aged 40-60 years old, n=8; O=old >65 years old, n=19) and in the North American (Canada) cohort (Y: n=3; M: n=4; O: n=6). **P** Frequencies of CD49f+-primitive HSC (defined as CD34+CD19-CD38-CD45RA-CD90+CD49f+ cells) within CD34+LIN- compartment in healthy donors from different ages in the North American (Canada) cohort (Y: n=3; M: n=4; O: n=6). Statistical test for correlation: Spearman r; Statistical test for multiple comparisons: Kruskal-Wallis test with Dunn's correction for multiple comparisons. (HSC, hematopoietic stem cells; MPP, multi-potent progenitors; MLP, multi-lymphoid progenitors; Pre-B/NK, B and NK precursors; ETP, early T progenitors; CMP, common myeloid progenitors; GMP, granulocyte-monocyte progenitors; MEP, megakaryocyte-erythrocyte progenitors; MKp, megakaryocyte progenitors, EP, erythrocyte progenitors).

### Supplementary Figure 4

**A**

| Confidence intervals for 1/(stem cell frequency) | | | | | | | | | |
|---|---|---|---|---|---|---|---|---|---|
| Group | 4 weeks | | | 12 weeks | | | 20 weeks | | |
| | Lower | Estimate | Upper | Lower | Estimate | Upper | Lower | Estimate | Upper |
| CB | 4328 | 1647 | 626 | 4409 | 1694 | 651 | 2158 | 504 | 118 |
| Young BM | 23582 | 11944 | 6050 | 19743 | 10052 | 5118 | 82481 | 36280 | 15958 |
| Middle age BM | 9207 | 4487 | 2187 | 2832 | 1208 | 515 | 37792 | 14537 | 5592 |
| Old BM | 10792 | 5574 | 2879 | 2419 | 1097 | 497 | 22988 | 12247 | 6525 |

| Pairwise tests for differences in stem cell frequencies | | | | | | | | |
|---|---|---|---|---|---|---|---|---|
| 4 weeks | | | 12 weeks | | | 20 weeks | | |
| Group 1 | Group 2 | P value | Group 1 | Group 2 | P value | Group 1 | Group 2 | P value |
| Middle age BM | Old BM | 0.666 | Young BM | Mid-age BM | 8.95E-05 | Young BM | Mid-age BM | 0.151 |
| Middle age BM | Bmyoung | 0.0579 | Young BM | Old BM | 2.73E-05 | Young BM | Old BM | 0.0197 |
| Middle age BM | CB | 0.0722 | Young BM | CB | 0.00437 | Young BM | CB | 7.03E-07 |
| Old BM | Bmyoung | 0.112 | Middle age BM | Old BM | 0.87 | Middle age BM | Old BM | 0.771 |
| Old BM | CB | 0.0226 | Middle age BM | CB | 0.621 | Middle age BM | CB | 0.000438 |
| Young BM | CB | 0.000355 | Old BM | CB | 0.514 | Old BM | CB | 0.000224 |

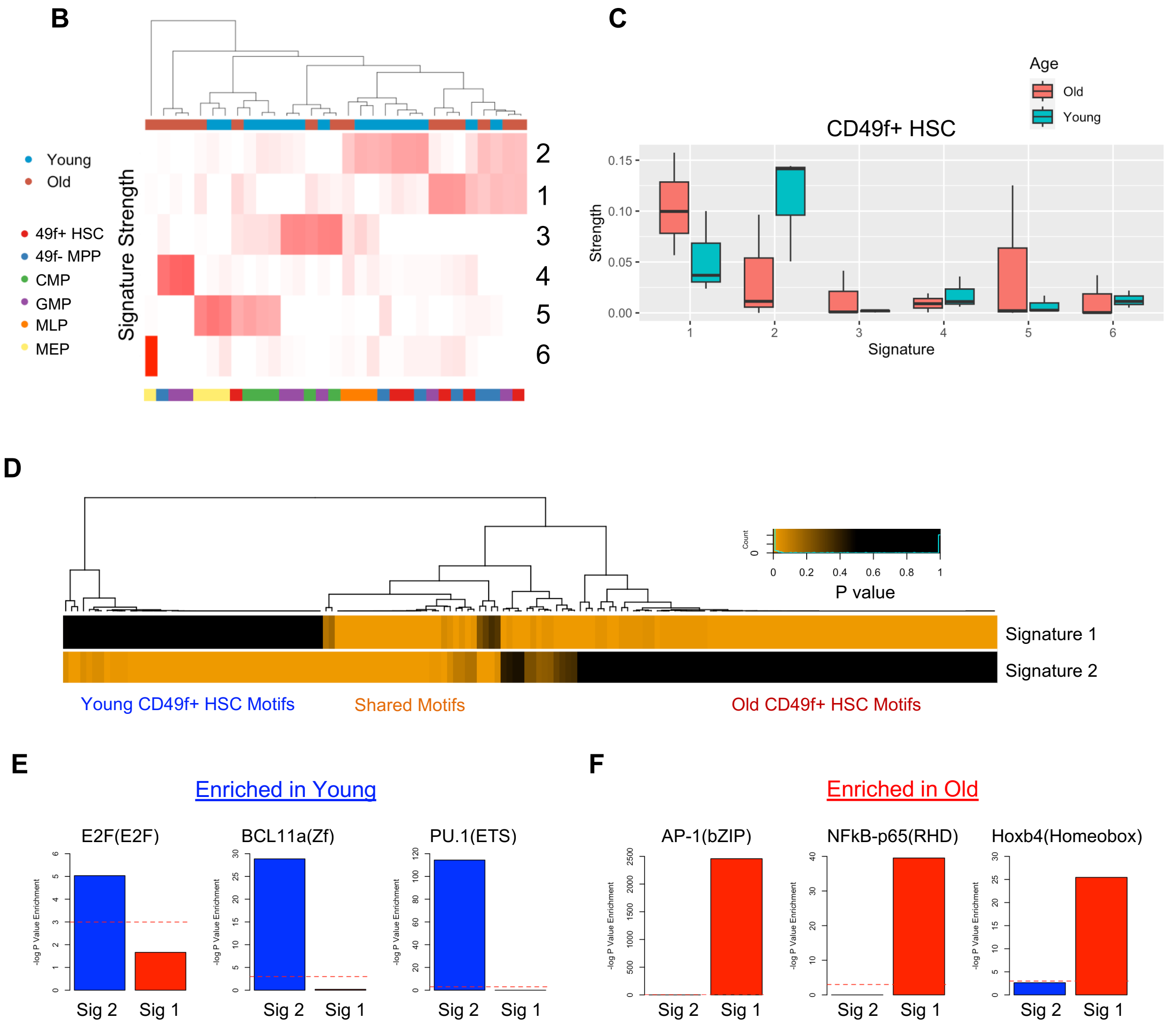


**Supplementary Figure 4. A.** Tables showing the confidence intervals for 1/stem cell frequency and p values from pairwise test for differences in the stem cell frequencies for the indicated sample groups as calculated by ELDA. Referring to Figure 2. **B.** Heatmap showing the signature strength of the six ATAC-seq signatures from NMF analysis of indicated BM HSPC subpopulations from young (3 donors, 21-28y, n=16 subpopulations) and old (4 donors, 52-79y, n=15 subpopulations). **C.** Signature strength assessment in CD49f+ HSC from young compared to old donors for 6 NMF signatures. **D.** Heatmap of the transcription factor motif enrichment of signature 1 and signature 2 shows three main clusters: enriched in signature 1 only (old CD49f+ HSC motifs), shared motifs and signature 2 only motifs (young CD49f+ HSC motifs). **E.** Log p value enrichment of indicated motifs enriched in young CD49f+HSC. Dotted red lines mark -log(0.05), indicating statistical significance. **F.** Log p value enrichment of indicated motifs enriched in old CD49f+HSC. Dotted red lines mark -log(0.05), indicating statistical significance.

#### Supplementary Figure 5

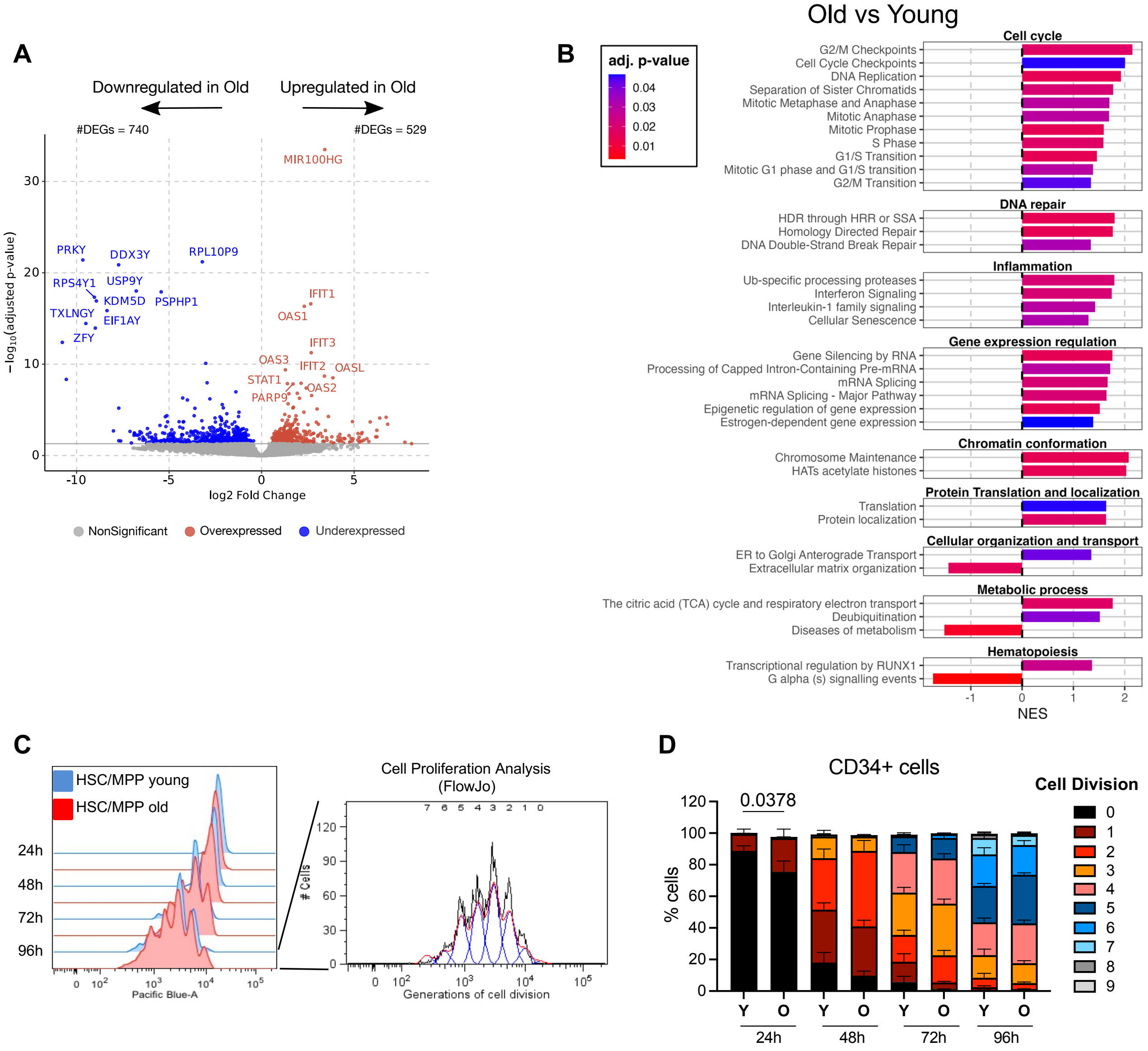


**Supplementary Figure 5. A.** Volcano plot showing significant (FDR < 0.05) downregulated and upregulated genes between old and young dataset after an overnight activation in culture. **B.** Bar-plot displaying significant (FDR < 0.05) GSEA categories from the Reactome Database used for the generation of the heatmap shown in Figure 3B. Bar length represents the normalized enrichment score (NES) for each category and the bar color shows the adjusted p-value (FDR= False Discovery Rate). **C.** (Left) Representative overlaid histogram in of the dilution of Cell Tracker dye during proliferation of young and aged sorted HSC+MPP populations at different time points. (Right) Automatic cell proliferation analyses performed by FlowJo software. **D.** Stacked bar graphs showing the frequencies of cells that completed one or more cell division overtime in culture in young (Y) and old (O) total CD34+ cells. Data are shown as Mean +/- SEM. Statistical test for comparison: Mann-Whitney.

### Supplementary Figure 6

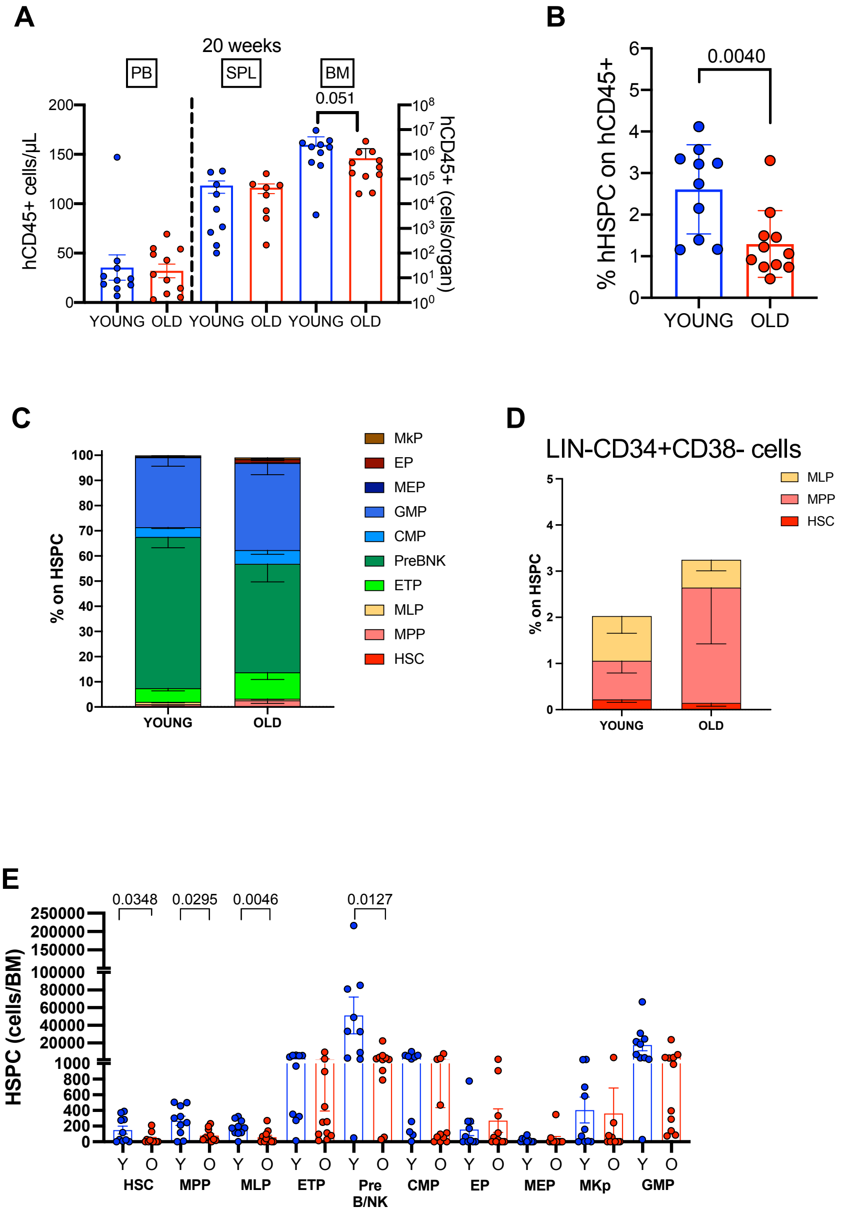


**Supplementary Figure 6. A.** Absolute quantification of human CD45+ cells in PB (plotted on the left axis), SPL and BM (plotted on the left axis) at 20 weeks after transplantation in mice transplanted with young and aged HSPCs. **B.** Percentage of human HSPC in the BM of mice transplanted with young or aged HSPCs at 20 weeks after transplantation. **C.** Composition of human BM HSPC compartment in mice transplanted with young or aged HSPCs at 20 weeks after transplantation. **D.** Percentage of human LIN-CD3+CD38- subpopulations within HSPC compartment in the BM of mice transplanted with young or aged HSPCs at 20 weeks after transplantation. **E.** Absolute quantification of human HSPC subpopulations in the BM of mice transplanted with young or aged HSPCs at euthanasia. In all the graphs data are shown as mean ± SEM.

### Supplementary Figure 7

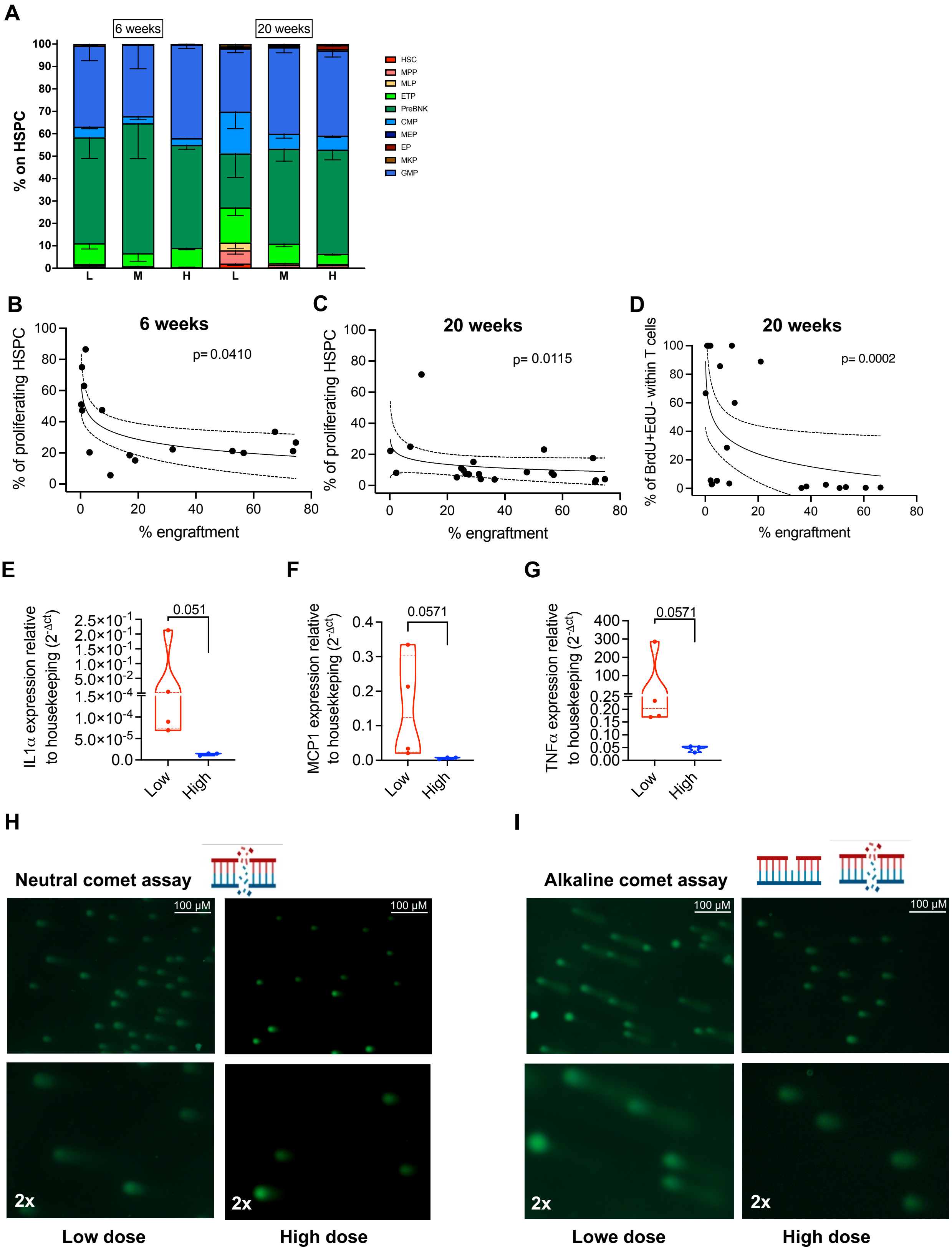

**Supplementary Figure 7. A.** Composition of human BM HSPC compartment in Low, Middle and High engrafted mice at both 6 and 20 weeks after transplantation. **B-C.** Correlation between the percentage of total proliferating HSPCs (BrdU+EdU-; BrdU+EdU+; BrdU-EdU+) with human engraftment in murine BM at 6 and 20 weeks after transplantation. **D.** Correlation between the percentage of T cells that entered S phase per hour (BrdU+EdU-) with human engraftment in PB at 20 weeks after transplantation**. E-G.** Relative mRNA expression of inflammatory genes *IL1*$\alpha$, *MCP1* and *TNF*$\alpha$ in human CD34+ cells isolated from Low and High engrafted mice at 20 weeks. **H-I.** Representative images of CD34+ isolated from Low (left) or High (right) engrafted mice analyzed by (**H**) neutral or alkaline (**I**) comet assays.
